# Convergent Evolution of Pain-Inducing Defensive Venom Components in Spitting Cobras

**DOI:** 10.1101/2020.07.08.192443

**Authors:** T.D. Kazandjian, D. Petras, S.D. Robinson, J. van Thiel, H.W. Greene, K. Arbuckle, A. Barlow, D.A. Carter, R.M. Wouters, G. Whiteley, S.C. Wagstaff, A.S. Arias, L-O. Albulescu, A. von Plettenberg Laing, C. Hall, A. Heap, S. Penrhyn-Lowe, C.V. McCabe, S. Ainsworth, R.R. da Silva, P.C. Dorrestein, M.K. Richardson, J.M. Gutiérrez, J.J. Calvete, R.A. Harrison, I. Vetter, E.A.B. Undheim, W. Wüster, N.R. Casewell

## Abstract

Convergent evolution provides unparalleled insights into the selective drivers underlying evolutionary change. While snakes use venom primarily for predation, and venom composition often reflects diet specificity, three lineages of spitting cobras have independently evolved the ability to use venom as a defensive projectile. Using gene, protein and functional analyses, we show that the three spitting lineages possess venom characterized by an upregulation of PLA_2_ toxins, which potentiate the action of venom cytotoxins to activate mammalian sensory neurons and cause enhanced pain. These repeated independent changes provide a fascinating example of convergent evolution across multiple phenotypic levels driven by exaptations. Notably, the timing of their origins suggests that defensive venom spitting may have evolved in response to the emergence of bipedal hominids in Africa and Asia.

**One Sentence Summary:** Venom spitting by snakes coincides with the emergence of hominins and is underpinned by convergent increases in pain-enhancing toxins

## Main Text

Convergent evolution, the independent emergence of similar traits, provides unparalleled opportunities to understand the selective drivers and mechanisms of adaptation (*1*). Thanks to their discrete function and direct genotype-phenotype link, animal venoms represent a superlative model system for unravelling both the driving forces and the underlying genetic mechanisms of molecular adaptation. Snake venoms consist of variable mixtures of proteinaceous components that cause potent hemotoxic, neurotoxic and/or cytotoxic pathologies in both prey and potential adversaries, including humans (*2*). Previous evidence suggests that venom variation is largely driven by dietary variation (*3*), but defensive drivers of snake venom evolution are rarely considered (although see (*4*)). The evolution of venom projection or ‘spitting’ in cobras offers an excellent model system for exploring the evolution of defensive toxins, as this behavior has no role in prey capture, targets specific sensory tissues, and is the only long-distance, injurious defensive adaptation among almost four thousand species of snakes. Remarkably, venom spitting evolved independently three times, all within a single clade of closely related elapid snakes (*5, 6*): the African spitting cobras (*Naja*: subgenus *Afronaja*), Asian spitting cobras (*Naja*: subgenus *Naja*) and the rinkhals (monotypic *Hemachatus*). All use modified fangs with small, rounded, front-facing orifices (*7*) to produce a spray of venom, which may reach distances of 2.5 m (*8*), and targets the eyes of an aggressor (*9*) (Fig. S1). This suite of behavioral, morphological, and biochemical traits results in intense ocular pain and inflammation, which can lead to the loss of eyesight (*10*). The three independent origins of spitting, solely within a clade more generally characterized by the visually defensive behavior of hooding (*5, 6*), provides an ideal model system to test whether similar selective pressures have resulted in convergent changes in molecular venom composition and overlying morphological and behavioral levels.

We used a multi-disciplinary approach consisting of transcriptomic, proteomic, functional and phylogenetic comparisons of 17 widely distributed elapids: 14 *Naja* (true cobras), the rinkhals *Hemachatus haemachatus*, and two non-spitting immediate outgroup species, *Walterinnesia aegyptia* and *Aspidelaps scutatus*, to investigate the evolutionary consequences of venom spitting. First, we reconstructed the phylogeny of these snakes using a multilocus coalescent species tree approach based on sequences from two mitochondrial and five nuclear genes. Fossil-calibrated molecular dating suggests that spitting originated in African spitting cobras 6.7-10.7 million years ago (MYA), and likely around 4 million years later in the Asian spitting cobras (2.5-4.2 MYA) (Fig. 1A). The origin of spitting in *Hemachatus* could not be dated, other than determining that it occurred <17 MYA, following its divergence from true cobras (*Naja*) (Fig. 1A).

**Fig. 1.**
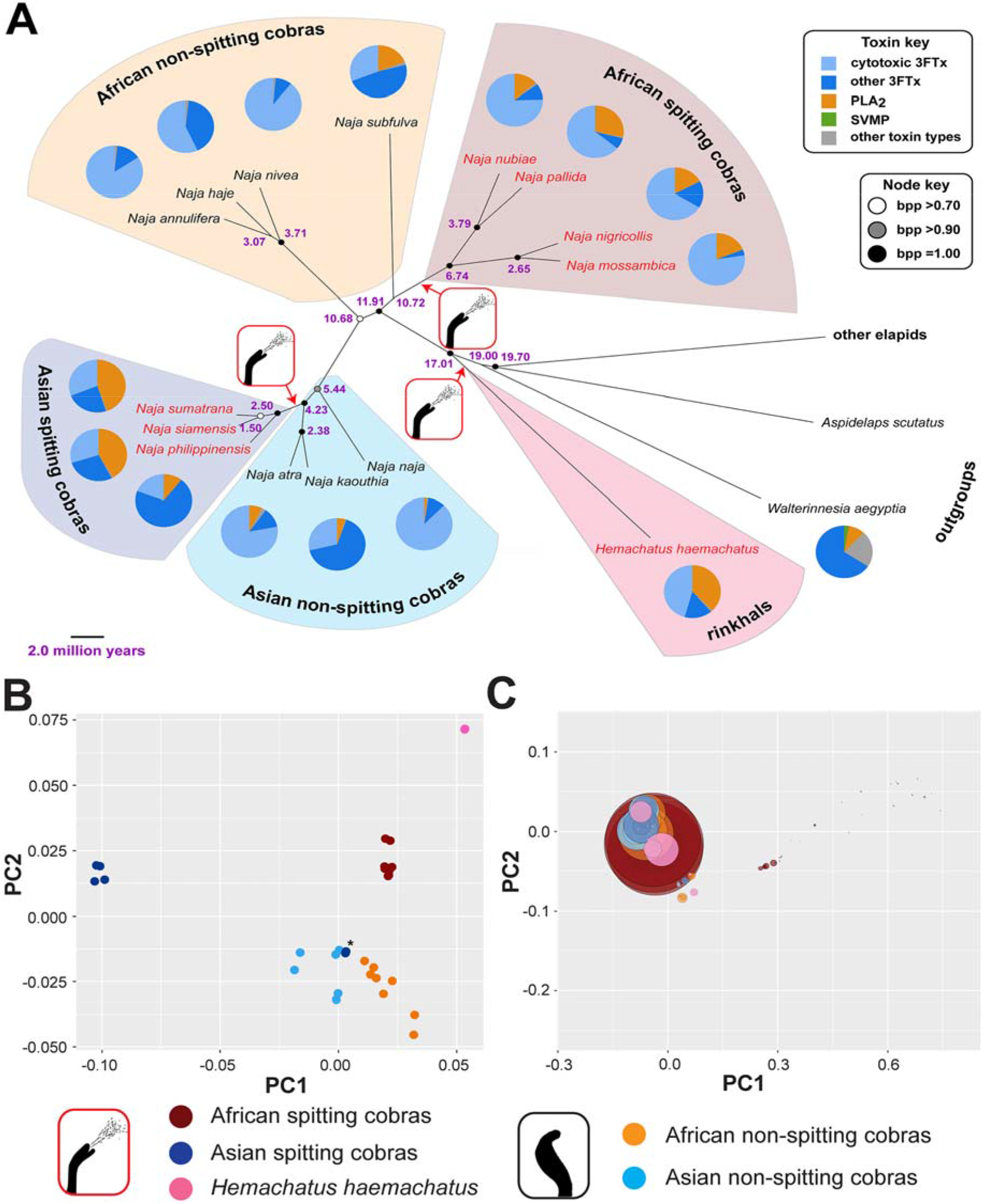
Reconstruction of the evolutionary origin of venom spitting and comparative analysis of venom composition. (A) The species tree is derived from five nuclear and two mitochondrial markers reconstructed using a multispecies coalescent model in *BEAST and pruned to display the taxa whose venoms were analyzed in this study. Support for the various nodes is indicated by colored circles, representing the following Bayesian posterior probabilities (bpp): black = 1.00, grey >0.90, white >0.70. Purple node labels indicate calculated divergence times from the *BEAST analysis (see Fig. S13 for calculated divergence time ranges). Spitting species are highlighted by red tip labels, and the three independent origins of venom spitting are indicated by the red-boxed spitting images and arrows. Pie charts adjacent to the tip labels represent the proteomic toxin composition of each of the sampled species as a percentage of total toxin proteins: light blue, cytotoxic 3FTx; dark blue, other 3FTx; orange, PLA_2_; green, SVMP; grey, all other toxin types. (B) Principal Coordinate Analysis (PCoA) of cobra (*Naja* spp.) and rinkhals (*H. haemachatus*) venom toxins reveal major distinctions between the spitting (red, Africa; dark blue, Asia; pink, *H. haemachatus*) and non-spitting (light blue, Asia; orange, Africa) lineages. The asterisk highlights Asian spitting species *N. philippinensis*, which exhibits greater similarity to non-spitting Asian and African species than to its nearest relatives (other Asian spitting cobras). Note each species is represented by two, typically overlapping, data points, which represent technical proteomic duplicates. (C) PCoA of cobra (*Naja* spp.) and rinkhals (*H. haemachatus*) CTXs reveals that the most abundant CTXs detected in venom exhibit little sequence diversity among spitting and non-spitting lineages. PCoA analysis was performed on a Euclidean distance matrix containing CTX amino acid data derived from top-down venom proteomics. Circle sizes reflect relative abundances of CTXs detected in the venom proteomes. The different lineages are colored as in B.

Next, we used a top-down proteomics approach underpinned by venom gland transcriptomic data (*11*) to characterize the toxins found in the venom of each species. Those analyses demonstrated that all cobra venoms are dominated by three finger toxins (3FTX), while in many species phospholipases A2 (PLA2) are the second most abundant toxin family (Fig. 1A, Fig. S2). Principal coordinate analysis (PCoA, Bray-Curtis), performed on a proteomic data-derived venom composition matrix of the ingroup species (*Naja* & *Hemachatus* spp.), separated the spitting lineages into three clusters that are distinct from the homogeneous cluster of venoms from non-spitters (Fig. 1B). The sole exception to this was *N. philippinensis*, a species reported to have a purely neurotoxic venom despite being able to spit (*12*), and whose venom composition placed it alongside non-spitting species (see Fig. 1B). Nonetheless, these findings demonstrate that each spitting cobra lineage exhibits distinct venom compositions that collectively differ from those of non-spitting cobras – a finding consistent with differences in venom-induced pathology observed following bites to humans (*13*).

3FTXs are major venom components in many elapid snakes (*14*) (Table S1). They are encoded by a multilocus gene family, resulting in numerous functionally distinct isoforms, including neurotoxins that interfere with various neuromuscular receptors, and cytotoxins that disrupt cell membranes to cause cytotoxicity (*15*). Proteomic data revealed that cytotoxic 3FTXs (CTXs) are typically the most abundant toxins found in *Naja* and *Hemachatus* venoms (mean 57.7% of all toxins), contrasting with the sister group species *W. aegyptia* and *A. scutatus* (*16*) and other elapids (Fig. 1A, Figs. S2, S3, Table S1). Despite ocular cytotoxicity presumably stimulating pain of value for enemy deterrence, we found no significant difference in the abundances of CTXs between spitting and non-spitting species (PGLS; t = -0.83, df = 15, p = 0.42) (Table S2), and ancestral state estimations suggest that the origin of CTX-rich venom preceded that of venom spitting (Fig. S4) (*5*). Moreover, PCoA analysis of a CTX Euclidean distance matrix derived from venom proteomic data revealed that all highly abundant cobra CTXs cluster tightly together (Fig. 1C), regardless of spitting ability. These findings extended to functional activities: measures of irritation, stimulated by high doses (1 mg/mL) of cobra venoms applied topically to non-sentient chick embryos (Tables S3, S4), revealed no association between cytotoxicity (Figs. S5, S6) and venom spitting (PGLS; t = 1.08, df = 15, p = 0.30) – results consistent with prior reports of comparable cytotoxicity to mammalian cells across the venoms of cobra species (*5*). Thus, the main toxins thought to be responsible for cytotoxic venom effects are similar across the different spitting and non-spitting cobra species.

Pain inflicted via the slow development of cytotoxicity may be less defensively relevant than that caused by a more immediate route, that of direct, algesic activity. To investigate venom-induced nociception, we assessed, via calcium imaging, the activation of trigeminal neurons – sensory neurons derived from the trigeminal ganglia that innervate the face and eyes (Fig. S7). Venoms from all cobra species activated trigeminal sensory neurons, though a lack of substantial activity was notable in African non-spitting cobras and the Asian non-spitting cobra *Naja kaouthia* (Fig. S7). The mechanism of activation for all venoms was characterized by rapid noncell-type-specific calcium influx followed by dye release from the cells into the extracellular space, indicative of non-specific disruption of cell membranes. We next determined potency values for each venom in sensory neuron-derived F11 cells. The resulting half maximal effective concentrations (EC50) demonstrated that spitting cobras had more potent venoms (PGLS; t = -4.48, df = 15, p = 0.0007) (Fig. 2A, Fig. S8, Table S2). These data therefore strongly support the hypothesis that the emergence of venom spitting in cobras is associated with convergent elevations in venom-induced activation of sensory neurons, and that spitting cobra venoms are more effective in causing pain than their non-spitting counterparts.

**Fig. 2.**
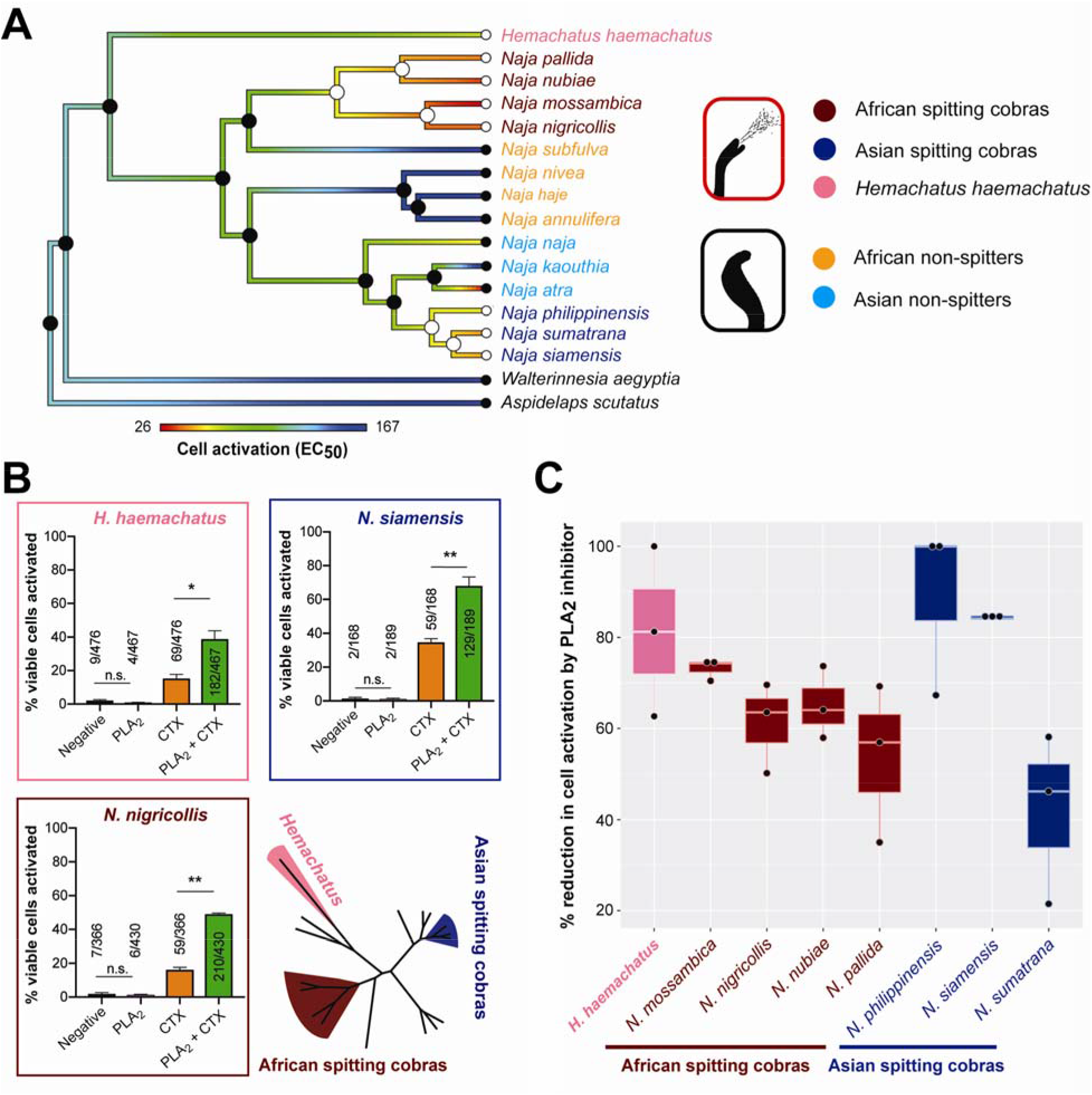
Spitting cobra venoms cause significantly greater activation of sensory neurons than non-spitting cobras, mediated via potentiation by PLA2 toxins. (A) Ancestral state estimation of the half maximal effective concentrations (EC_50_) of venom-induced activation of neuronal cells shows a significant association between venom potency and venom spitting (PGLS, t = -4.48, p = 0.0004). EC_50_ values were derived from concentration-response curves fitted to the data using a four-parameter Hill equation with variable Hill slope, and are expressed as the mean of triplicate measurements. Colored branches are scaled according to levels of cell activation (red, low EC50 and thus high venom potency; blue, high EC50 and thus lower venom potency). At each node and tip of the tree, filled or empty circles represent estimated ancestral states of non-spitting or spitting, respectively, and colored tip labels correspond to different lineages detailed to the right: spitting (red, Africa; dark blue, Asia; pink, *H. haemachatus*) and non-spitting (light blue, Asia; orange, Africa). (B) PLA2 toxins in spitting cobra venoms potentiate the activating effect of CTXs on sensory neurons. A CTX fraction from each venom was added to dissociated mouse DRG neurons in the presence or absence of a corresponding PLA_2_ fraction (added 1 min prior). Neuronal activation (i.e. a rapid increase in [Ca^2+^]_i_) was monitored following the first (1 min window) and second (2 min window) additions and presented as a proportion of viable cells. Data are expressed as mean ± SEM and are representative of 2-3 experiments. Statistical comparisons were made using unpaired parametric *t*-tests in Graphpad Prism (version 8.02). *, p < 0.05; **, p < 0.01. (C) The PLA_2_ inhibitor varespladib reduces neuronal activation stimulated by spitting cobra venoms. Calcium influx in F11 cells was measured on a FLIPR instrument incubated in the presence of venom from spitting species (2.4 μg or 4.8 μg [in the case of *H. haemachatus* and *N. philippinensis*] venom) and in the presence or absence of varespladib (13 μM). The data displayed represents the percentage of venom only cell activation stimulated by treatment with venom and varespladib. Values are max-min (n = 3). The error bars represent standard error of the mean (SEM) with the box mid-lines representing the median value.

To determine the toxins responsible for this effect, we repeated these experiments using fractionated venom from three representative spitting cobras (*N. nigricollis*, African; *N. siamensis*, Asian; *H. haemachatus*, rinkhals) (Fig. S9). For all species tested, only fractions corresponding to CTXs activated sensory neurons, while those corresponding to other toxins (e.g. neurotoxins, PLA_2_s, etc) were inactive (Fig. S9). However, none of the CTX fractions by themselves completely recapitulated the effects of whole venom or those of re-pooled venom fractions, suggesting that multiple venom components are acting synergistically.

PLA_2_s are nearly ubiquitous, typically enzymatic, multifunctional toxin components found in snake venoms (*14, 15*). As the hemolytic activity of CTXs was previously shown to be potentiated by PLA2 toxins (*17*), we hypothesized that, although venom PLA2s are not independently capable of activating sensory neurons, they may potentiate the effects induced by CTXs. Consequently, for each of the three representative spitting cobra venoms, we quantified the activation of sensory neurons stimulated by CTX fractions in the presence or absence of a corresponding PLA_2_ venom fraction. Consistent with our expectation, the proportion of viable sensory neurons activated by each CTX fraction was significantly increased when preceded by the addition of a PLA_2_ fraction, and this result was consistent for each representative of the three spitting lineages (unpaired t-test; *N. nigricollis*, t = 18.77, df = 2, p = 0.003; *N. siamensis*, t = 5.75, df = 4, p = 0.005; *H. haemachatus*, t = 4.18, df = 4, p = 0.01) (Fig. 2B, Fig. S10). To confirm the pain-potentiating activity of PLA_2_ toxins, we compared the potency of sensory neuron activation caused by whole venoms from our eight spitting species in the presence or absence of the PLA_2_ inhibitor varespladib. Significant reductions in sensory neuron activation occurred in the presence of this inhibitor (unpaired t-test; t = 2.77, df = 14, p = 0.02) (Fig. 2C, Fig. S11), providing further compelling evidence that PLA_2_s potentiate CTX effects on sensory neurons.

In line with the above findings, comparative analysis of (i) the proteomic abundance of PLA_2_ toxins and (ii) the quantified enzymatic PLA2 activity, determined via specific in vitro colorimetric assay, revealed that spitting cobra venoms have significantly higher PLA2 abundance and activity than those of non-spitting species (PGLS; t = 4.24, df = 15, p = 0.0007 and t = 2.24, df = 15, p = 0.04, respectively) (Fig. 3A, 3B, Fig. S12, Table S2). Analysis of a PLA2 Euclidean distance matrix derived from venom proteomic data revealed substantial distinctions between the different lineages of spitting and non-spitting cobras, particularly in the case of African species (Fig. 3C). Additionally, *H. haemachatus* PLA_2_s clustered tightly with those from African spitting cobras (Fig. 3C), despite the divergence of *Hemachatus* ~17 MYA. These findings suggest that the PLA_2_s of African spitting cobras and *H. haemachatus* exhibit some evidence of molecular convergence, and divergence from the PLA_2_s of Asian spitting cobras (Fig. 3C). The combination of these data, plus evidence from the sensory neuron assays, demonstrate that convergent evolution of spitting is tightly linked with convergent upregulation of PLA_2_ toxins in those venoms.

**Fig. 3.**
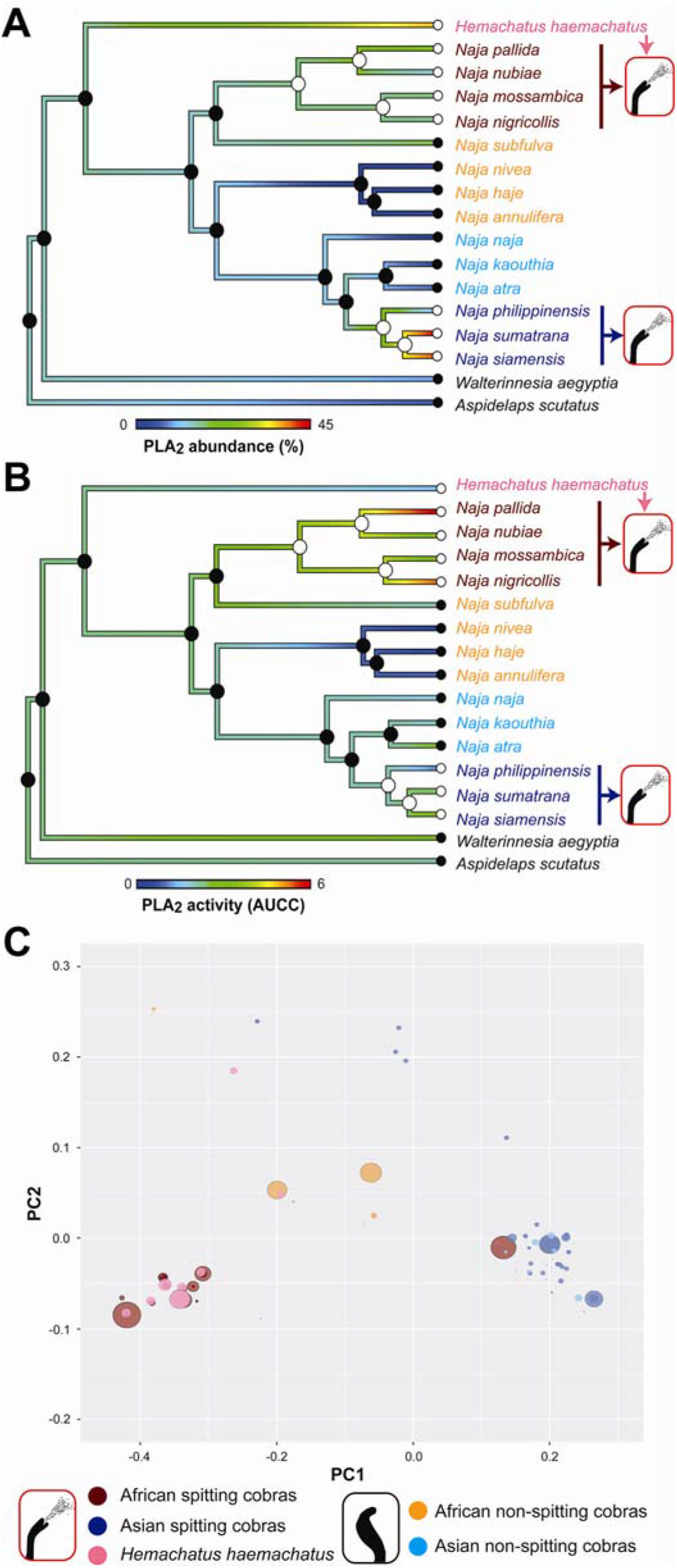
The abundance, enzymatic activity and diversity of phospholipase A_2_ (PLA_2_) toxins is associated with convergent evolution of venom spitting. (A) Ancestral state estimation of proteomic abundance of PLA_2_s, represented as the percentage of PLA_2_ toxins in the venom proteomes, revealed a significant association with venom spitting (PGLS, t = 4.24, p = 0.0007). Colored branches are scaled according to PLA2 abundance (blue, low abundance; red, high abundance). At each node and tip of the tree, filled or empty circles represent the estimated ancestral state of non-spitting or spitting, respectively, and colored tip labels correspond to the different lineages detailed in C. (B) Ancestral state estimation of enzymatic PLA_2_ activities, determined via area under the curve of concentration curves (AUCC) analysis of a kinetic *in vitro* colorimetric assay, revealed a significant association with venom spitting (PGLS, t = 2.24, p = 0.04). Colored branches are scaled according to PLA_2_ activity (blue, low activity; red, high activity). Labels as in (A), and see Fig. S12 for PLA_2_ activity concentration curves. (C) Principal Coordinate Analysis (PCoA) of cobra (*Naja* spp.) and rinkhals (*H. haemachatus*) PLA_2_ toxins reveals major variation between African spitting and non-spitting lineages, but little variation between Asian spitting and non-spitting lineages. Note also the convergent placement of *Hemachatus* PLA_2_ toxins with those of African spitting cobras. PCoA analysis was performed on a Euclidean distance matrix containing PLA_2_ amino acid data derived from top-down venom proteomics. Circle sizes reflect relative abundances of PLA_2_s detected in the venom proteomes. Spitting lineages are colored red (African), dark blue (Asian) and pink (*H. haemachatus*) and non-spitting lineages are colored light orange (African) and light blue (Asian).

To exclude the possibility that functional distinctions simply reflect general differences in venom potency, e.g., venom lethality for prey capture, rather than specifically defensive adaptations, we tested each of the venoms in a murine lethality assay (median lethal dose, LD_50_). These experiments revealed highly variable venom potencies (2.36-33.75 μg/mouse across *Naja*) (Table S5) and no significant differences between spitting and non-spitting species (PGLS; t = 0.86, df = 15, p = 0.40) (Table S2). This strongly suggests that the enhanced pain caused by spitting cobras is explicitly associated with the defensive use of those venoms, rather than being an evolutionary spandrel from selection for prey subjugation.

While the Asian cobras *N. atra* and *N. kaouthia* have been considered as non-spitters throughout this study, recent reports suggest that certain individuals/populations of these two species show some spitting ability (*18, 19*). However, reanalysis of our data with *N. atra* and *N. kaouthia* treated as spitting species revealed only minor differences in statistical significance for the various analyses (Table S6). Importantly, our key results concerning associations between venom spitting and (i) increased PLA_2_ abundance and (ii) enhanced activation of sensory neurons remaining highly significant (PGLS: t = 2.86, df = 15, p = 0.01 and t = -4.59, df = 15, p = 0.0004, respectively; Table S6), suggesting that our choice of coding of these two species does not impact our conclusions.

Our results detail the genetic and functional consequences of the evolution of venom spitting and demonstrate that defense can be a major driver of snake venom composition. Spitting probably only evolved within one relatively small clade of Afro-Asian elapids due to the integrated exaptation of a unique combination of pre-existing behaviors and cytotoxic venom activities. Early evolution of cytotoxic venom activity in cobras and near relatives (~26 MYA; Fig. S4) has previously been linked to defense, as cytotoxicity co-originates with ‘hooding’ (*5*), a long-distance visual defensive display. This elevated posture, coupled with pre-emptive striking and accompanied by occasional premature releases of venom, may have provided a behavioral precursor for the evolution of more targeted venom spitting. Pre-existing CTXs, largely absent from the other elapid venoms (Table S1), likely provided the necessary baseline ocular toxicity to favor the inception and retention of spitting (*19*). Subsequent independent increases in PLA_2_ toxins, which act in synergy with pre-existing CTXs, resulted in increased venom-induced activation of nociceptors (Fig. 2). This potentiating effect of PLA_2_s may be crucial for causing immediate pain with sufficient intensity to rapidly deter aggressors, allowing the snake to escape. Consequently, our results from spitting cobras highlight how convergence at molecular, morphological, behavioral, and functional levels, in combination with behavioral and molecular exaptations, can drive complex adaptations in ecologically important traits.

Rare but repeatedly evolved adaptations likely result from similar but unexpected ecological relationships. Most discussions of defensive behavior involve potential predators, and certain mammals and birds commonly eat snakes (*20, 21*). However, predation events involving spitting cobras still occur often, therefore these interactions appear unremarkable in terms of evolutionary history and biogeography. Beyond predators, potential threats to snakes include inadvertent trampling and pre-emptive defensive killing. That spitting evolved to prevent snakes being trampled by herds of ungulates in African savannas (*22*) does not explain the existence of primarily forest-dwelling Asian spitting cobras (*23*). Moreover, large ungulates typically have laterally-located eyes and primarily use olfaction and hearing to detect danger, making them unlikely to be especially vulnerable to spitting.

Several considerations make ancient hominins a more compelling candidate for favoring repeated evolution of spitting in the Afro-Asian cobras: (i) growing evidence suggests that snakes have profoundly influenced primate neurobiology and behavior (*24*), and that interactions between these two lineages have been important, sometimes reciprocally, throughout the 75 million year history of primates (*25, 26*). (ii) Compared to obligately carnivorous mammals, anthropoid primates are more visually acute, cognitively complex, and culturally sophisticated (*24*). (iii) Diverse anthropoids mob snakes, and some distinguish between harmless and dangerous species, eating the former and killing the latter with clubs or projectiles (*25–27*). (iv) Characteristics (ii) and (iii) are enhanced among bipedal, larger brained hominins (*24, 26*), which could have posed a singularly important, long-distance threat to dangerous Afro-Asian snakes (*28*). The initial divergence of Africa spitting cobras as recently as 6.7 MYA (Fig. 1) occurred soon after the divergence of hominins from *Pan* (bonobos and chimpanzees) ~7 MYA, thus coinciding with the early evolution of bipedalism, enlarged brains, tool use, and occupation of savanna habitats by the former (*29*). Likewise, the origin of the Asian spitter clade ~2.5 MYA is approximately contemporaneous with the arrival in Asia of *Homo erectus* (*30, 31*). Additional fossils and more finely-tuned dating of relevant cobra and primate divergences will enable further testing of this hypothesis. Nonetheless, the repeated evolution of spitting cobras might well amount to an expansion of ‘Lucy’s Legacy’ (*32*), alluding to the famous *Australopithecus afarensis* fossil that initially shed so much light on bipedalism and big brains in our own lineage.

## Acknowledgments

The authors thank Paul Rowley and Edouard Crittenden for maintenance of snakes and performing venom extractions, and Wendy Grail for technical support relating to the generation of the species tree. The NVIDIA TITAN-X GPU used for BEAST analyses was kindly donated by the NVIDIA Corporation.

## Funding

This work was funded from a studentship supported by Elizabeth Artin Kazandjian to T.D.K., grant PE 2600/1 from the German Research Foundation (DFG) to D.P., grant OPUS 1354156 from the US National Science Foundation to H.W.G., grants FAPESP 2017/18922-2 and 2019/05026-4 from the São Paulo Research Foundation to R.R.d.S, grants RPG-2012-627 and RFG-10193 from the Leverhulme Trust to R.A.H. and W.W., grant MR/L01839X/1 from the UK Medical Research Council to J.M.G., R.A.H., J.J.C. and N.R.C., fellowship DE160101142 from the Australian Research Council, and fellowship FRIPRO-YRT #287462 and grant DP160104025 from the Research Council of Norway to E.A.B.U., and a Sir Henry Dale Fellowship (200517/Z/16/Z) jointly funded by the Wellcome Trust and Royal Society to N.R.C.

## Author contributions

R.A.H., W.W. and N.R.C. conceived the research. T.D.K., D.P., K.A., M.K.R., R.A.H., J.J.C., I.V., E.A.B.U., W.W. and N.R.C. designed the research. T.D.K., D.P., S.R., J.vT., A.B., D.A.C., R.M.W., G.W., S.C.W., A.S.A., L-O.A., A.vP.L., C.H., A.H., S.P-L., C.V.M., S.A., R.R.d.S., P.C.D., J.M.G., J.J.C., I.V., E.A.B.U., W.W. and N.R.C. performed the research. T.D.K. and N.R.C. wrote the manuscript with major input from D.P., S.D.R., H.W.G., I.V., E.A.B.U. and W.W. All authors discussed and commented on the manuscript.

## Competing interests

Authors declare no competing interests.

## Data and materials availability

The molecular data associated with species tree generation have been deposited to the nucleotide database of NCBI and the accession numbers are displayed in Table S7. The transcriptome data have been deposited in the SRA and TSA databases of NCBI and are associated with the BioProject accession number PRJA506018. Mass spectrometry data and database search results for top-down and bottom-up proteomic experiments are publicly available in the MassIVE repository under accession number MSV000081885 and in proteomXchange with accession number PXD008597.

## Supplementary Materials

Materials and Methods

Figures S1-S13

Tables S1-S10

Data S1-S3

Supplementary References

Supplementary Acknowledgments

## Notes

### Competing Interest Statement

The authors have declared no competing interest.

### Summary of Updates

This version of the manuscript remains unchanged from the previous version. The supplementary information has, however, been revised to contain an updated version of Figure S13, which displays the cobra (and near relatives) species tree. The figure legend of Figure S13 has also been updated accordingly.

## References

1. J. Losos, Improbable Destinies: How Predictable is Evolution? (Penguin Random House, 2017).

2. J. M. Gutiérrez, J. J. Calvete, A. G. Habib, R. A. Harrison, D. J. Williams, D. A. Warrell, Snakebite Envenoming. Nat. Rev. Dis. Prim. 3, 17063 (2017).

3. J. C. Daltry, W. Wüster, R. S. Thorpe, Diet and Snake Venom Evolution. Nature. 379, 537–540 (1996).

4. H. Ward-Smith, K. Arbuckle, A. Naude, W. Wüster, Fangs for the Memories ? A Survey of Pain in Snakebite Patients Does Not Support a Strong Role for Defense in the Evolution of Snake Venom Composition. Toxins (Basel). 12, 1–20 (2020).

5. N. Panagides, T. Jackson, M. Ikonomopoulou, K. Arbuckle, R. Pretzler, D. Yang, S. Ali, I. Koludarov, J. Dobson, B. Sanker, A. Asselin, R. Santana, I. Hendrikx, H. van der Ploeg, J. Tai-A-Pin, R. van den Bergh, H. Kerkkamp, F. Vonk, A. Naude, M. Strydom, L. Jacobsz, N. Dunstan, M. Jaeger, W. Hodgson, J. Miles, B. Fry, How the Cobra got its Flesh-Eating Venom: Cytotoxicity as a Defensive Innovation and its Co-Evolution with Hooding, Aposematic Marking, and Spitting. Toxins (Basel). 9, 103 (2017).

6. W. Wüster, S. Crookes, I. Ineich, Y. Mane, C. E. Pook, J. F. Trape, D. G. Broadley, The Phylogeny of Cobras Inferred from Mitochondrial DNA Sequences: Evolution of Venom Spitting and the Phylogeography of the African Spitting Cobras (Serpentes: Elapidae: *Naja nigricollis* Complex). Mol. Phylogenet. Evol. 45, 437–453 (2007).

7. C. M. Bogert, Dentitional Phenomena in Cobras and Other Elapids with Notes on Adaptive Modifications of Fangs. Bull. Am. Museum Nat. Hist. 81, 260–285 (1943).

8. S. Rasmussen, B. Young, H. Krimm, On the ‘Spitting’ Behaviour in Cobras (Serpentes: Elapidae). J. Zool. 237, 27–35 (1995).

9. B. A. Westhoff, G., Boetig, M., Bleckmann, H., Young, Target Tracking During Venom ‘Spitting’ by Cobras. J. Exp. Biol. 213, 1797–1802 (2010).

10. L. D. Warrell, D. A., Ormerod, Snake Venom Ophthalmia and Blindness Caused by the Spitting Cobra *(Naja nigricollis)* in Nigeria. Am. J. Trop. Med. Hyg. 25, 525–529 (1976).

11. S. Ainsworth, D. Petras, M. Engmark, R. D. Süssmuth, G. Whiteley, L. O. Albulescu, T. D. Kazandjian, S. C. Wagstaff, P. Rowley, W. Wüster, P. C. Dorrestein, A. S. Arias, J. M. Gutiérrez, R. A. Harrison, N. R. Casewell, J. J. Calvete, The Medical Threat of Mamba Envenoming in Sub-Saharan Africa Revealed by Genus-wide Analysis of Venom Composition, Toxicity and Antivenomics Profiling of Available Antivenoms. J. Proteomics. 172, 173–189 (2018).

12. G. Watt, L. Padre, M. L. Tuazon, R. D. G. Theakston, L. Laughlin, Bites by the Philippine Cobra (*Naja naja philippinensis*): Prominent Neurotoxicity with Minimal Local Signs. Am. J. Trop. Med. Hyg. 39, 306–311 (1988).

13. J. Meier, J. White, Handbook of Clinical Toxicology of Animal Venoms and Poisons (Informa Healthcare USA Inc, New York, ed. 1st, 2008).

14. T. Tasoulis, G. K. Isbister, A Review and Database of Snake Venom Proteomes. Toxins (Basel). 9, 290 (2017).

15. C. R. Ferraz, A. Arrahman, C. Xie, N. R. Casewell, R. J. Lewis, J. Kool, F. C. Cardoso, Multifunctional Toxins in Snake Venoms and Therapeutic Implications: From Pain to Hemorrhage and Necrosis. Front. Ecol. Evol. 7, 1–19 (2019).

16. G. Whiteley, N. R. Casewell, D. Pla, S. Quesada-Bernat, R. A. E. Logan, F. M. S. Bolton, S. C. Wagstaff, J. M. Gutiérrez, J. J. Calvete, R. A. Harrison, Defining the Pathogenic Threat of Envenoming by South African Shield-nosed and Coral Snakes (Genus *Aspidelaps),* and Revealing the Likely Efficacy of Available Antivenom. J. Proteomics. 198, 186–198 (2019).

17. E. Condrea, Membrane-active Polypeptides from Snake Venom: Cardiotoxins and Haemocytotoxins. Experientia. 30, 121–129 (1974).

18. A. Paterna, Spitting Behaviour in the Chinese Cobra *Naja atra*. Herpetol. Bull. (2019).

19. V. Santra, W. Wüster, *Naja kaouthia* (Monocled Cobra). Behaviour/Spitting. Herpetol. Rev. 48, 455–456 (2017).

20. H. W. Greene, in Biology of the Reptilia. Vol. 16: Ecology B: Defense and Life History, C. G. and R. B. Huey, Ed. (Alan R. Liss, Inc., New York, ed. 1st, 1988), pp. 1–152.

21. B. Van Valkenburgh, R. K. Wayne, Carnivores. Curr. Biol. 20, R915–R919 (2010).

22. T. Barbour, Rattlesnakes and Spitting Snakes. Copeia, 26–28 (1922).

23. E. R. Chu, S. A. Weinstein, J. White, D. A. Warrell, Venom Ophthalmia Caused by Venoms of Spitting Elapid and Other Snakes: Report of Ten Cases with Review of Epidemiology, Clinical Features, Pathophysiology and Management. Toxicon. 56, 259–272 (2010).

24. L. A. Isbell, The Fruit, the Tree, and the Serpent: Why We See So Well (Harvard University Press, Cambridge, 2009).

25. T. N. Headland, H. W. Greene, Hunters-gatherers and Other Primates as Prey, Predators, and Competitors of Snakes. Proc. Natl. Acad. Sci. 108, E1470–E1474 (2011).

26. H. W. Greene, Tracks and Shadows: Field Biology as Art (University of California Press, Berkeley and Los Angeles, 2013).

27. H. W. Greene, Evolutionary Scenarios and Primate Natural History. Am. Nat. 190, S69–S86 (2017).

28. S. Faurby, D. Silvestro, L. Werdelin, A. Antonelli, Brain Expansion in Early Hominins predicts carnivore extinctions in East Africa. Ecol. Lett. 3, 537–544 (2020).

29. L. Pozzi, J. A. Hodgson, A. S. Burrell, K. N. Sterner, R. L. Raaum, T. R. Disotell, Primate Phylogenetic Relationships and Divergence Dates Inferred from Complete Mitochondrial Genomes. Mol. Phylogenet. Evol. 75, 165–183 (2014).

30. S. Prat, First Hominin Settlements out of Africa. Tempo and Dispersal mode: Review and Perspectives. Comptes Rendus - Palevol. 17, 6–16 (2018).

31. F. Han, J. J. Bahain, C. Deng, É. Boëda, Y. Hou, G. Wei, W. Huang, T. Garcia, Q. Shao, C. He, C. Falguères, P. Voinchet, G. Yin, The Earliest Evidence of Hominid Settlement in China: Combined Electron Spin Resonance and Uranium Series (ESR/U-series) Dating of Mammalian Fossil Teeth from Longgupo Cave. Quat. Int. 434, 75–83 (2017).

32. A. Jolly, Lucy’s Legacy: Sex and Intelligence in Human Evolution (Harvard University Press, ed. 1st, 1999).

