## Supplementary material for "Convergent Evolution of Pain-Inducing Defensive Venom Components in Spitting Cobras": Fig. S1

**This PDF file includes:**

Materials and Methods

Figs. S1 to S13

Tables S1 to S10

Data Files S1 to S3

Supplementary References

Supplementary Acknowledgments

### Materials and Methods

#### Venoms and venom glands

Venoms from cobras and related species were pooled from wild-caught or captive bred (CB) specimens maintained in the Liverpool School of Tropical Medicine Herpetarium (Table S8). Crude venoms were lyophilized post-extraction and stored at 4°C until analysis. For venom gland transcriptomics, single specimens from each species were euthanized three days after venom extraction, and dissected venom glands were immediately flash frozen in liquid nitrogen and stored cryogenically until use. Details of the specimens used in this study are displayed in Table S8.

#### Species tree reconstruction

We reconstructed the phylogenetic relationships and divergence times of 46 species of cobra and related elapid taxa, representing almost all species of *Naja* and cobra-clade elapids under a multispecies coalescent model. This analysis was based on DNA sequence alignments of six genetic loci: mitochondrial DNA (partial CytB and ND4 gene sequences; Table S9); and the phased partial exon sequences of the nuclear genes CMOS, NT3, PRLR, UBN1 and RAG1. Briefly, we obtained blood samples, ventral scale clips, shed skins or other tissue in the field or from captive collections. Whole genomic DNA was extracted using a Qiagen DNeasy$^{TM}$ Tissue Kit (catalogue no. 69506 – www.qiagen.com), following the manufacturer's instructions, except for blood samples, where PBS buffer was not used and 200 µL of blood in Tris-EDTA buffer was added to 20 µL proteinase K. Where shed skins were used, the samples were left to lyse overnight. PCR was carried out in 15 µL volumes, using ThermoScientific 2X DreamTaq buffer. Primer sequences and thermocycling parameters are given in Table S9. Sanger dideoxy sequencing was carried out by Macrogen (dna.macrogen.com), using the forward primer for mitochondrial genes and forward and reverse primers for nuclear loci.

Sequences were checked, trimmed, translated into amino acid sequences and checked for unexpected indels or non-sense codons in CodonCode Aligner version 3.7.1 (www.codoncode.com). In nuclear locus sequences, heterozygous positions were replaced with the IUPAC uncertainty codes. The alleles of length heterozygotes were reconstructed using the online tool Indelligent 1.2 (http://dmitriev.speciesfile.org/indel.asp) (*33*). Individual allele sequences (haplotypes) were then estimated from diploid nuclear loci using the software PHASE v. 2.1.1 (*34*, *35*) over 5000 iterations with a burn-in of 500 and a thinning interval of 10, after preparation of the sequence data using SEQPHASE (*36*). PHASE was run three times to confirm burn-in and convergence across multiple runs. PHASE was run separately by genus, or, in the case of *Naja* by subgenus.

Details of the final DNA sequence alignments were as follows: mitochondrial DNA, 1,316 bp, 281 sequences; CMOS, 628 bp, 206 sequences; NT3, 657 bp, 340 sequences; PRLR, 551 bp, 456 sequences; UBN1, 504 bp, 414 sequences; RAG1, 879 bp, 281 sequences. Sample details are shown in Table S9 and NCBI nucleotide database accession numbers in Table S7. For phylogenetic reconstruction, we used a multispecies coalescent model employed in *BEAST in BEAST v.1.8.2 (*37*). Our approach was to initially generate a species tree for the entire set of 46 taxa, including virtually all cobras and cobra-like elapids, and then to extract the tree of the focal 17 species from that tree. The multispecies coalescent model implemented in *BEAST coestimates individual gene (locus) trees embedded within the overall species tree while accounting for incomplete lineage sorting. It requires prior assignment of sequences to species, and that each of these taxa be represented by at least one sequence of each locus. We lacked CMOS and RAG 1 sequences for certain taxa (*N. christyi* and *N. sputatrix*), so a single "dummy" sequence consisting of Ns was inserted for each of these taxa into the respective alignments in order to initiate the analysis.

Preliminary *BEAST runs using node calibrations based on the fossil record to estimate divergence times on an unconstrained species tree failed to converge, indicating overparameterisation. To overcome this, we adopted a two-step approach. First, an uncalibrated analysis was carried out estimating the topology of the species tree with the substitution rate of the mitochondrial alignment fixed at an arbitrary value and the substitution rate of the other gene alignments estimated relative to it. Then, a calibrated analysis was carried out to estimate divergence times by applying internal node calibrations based on the fossil record, with the monophyly of all species tree nodes that were well supported in the uncalibrated analysis (posterior probability > 0.99) constrained.

For the uncalibrated analysis, the precise model specification was determined by trialing more complex models initially, and then incrementally reducing model complexity (in terms of the number of free parameters) in order to achieve convergence. A Yule process was specified for the species tree with a piecewise constant population size model. A single GTR+G nucleotide substitution model with empirical base frequencies was specified for nuclear DNA, by linking the substitution model partitions of the respective five nuclear gene alignments. A separate GTR+G nucleotide substitution model with estimated base frequencies was specified for the mitochondrial alignment. A separate, unlinked strict molecular clock model was specified for each locus. The substitution rate of the mitochondrial locus was arbitrarily fixed to 1, and the substitution rates of the five nuclear loci estimated relative to it by assigning uninformative, uniform priors between 0 and 1. It was necessary to run the MCMC chain for hundreds of millions of generations, sampling every 1,000 generations, to achieve burn in and adequate sampling of parameters. This was run until the physical limits of the computer's hard drive were reached, at which time all parameters had burned in and achieved effective sample sizes > 200, except for two parameters which were > 170, assessed using the program Tracer. Trees sampled during burn in were removed, and the posterior sample of trees downsampled to approximately 50,000 trees using the program LogCombiner. The maximum clade credibility tree was then selected from these ~50,000 trees, with node heights centered on the median of the subsample, using TreeAnnotator.

For the calibrated analysis, the model specification was identical except the monophyly of species tree nodes that were well supported in the uncalibrated analysis were enforced, and node calibrations based on the fossil record were used to estimate divergence timings. These were: an exponential prior with a 0.01 MY mean and 17.0 MY offset on the MRCA of *Naja*, *Hemachatus* and *Pseudohaje*, which effectively fixed the age of this node at 17 MYA (*38*); since fossil calibrations constrain the minimum but not the maximum age of a node, this represents the youngest plausible age of this node. We also applied a uniform prior between 0 and 10 MY on the MRCA of the Australian genera *Oxyuranus* and *Pseudechis*, effectively placing a maximum divergence time on these genera of 10 MYA (*38*). The substitution rate of the mitochondrial locus was estimated based on these calibrations within an uninformative, uniform prior between 0 and 2% per million years. The substitution rates of the nuclear loci were estimated using uninformative uniform priors between 0 and 1.5% per million years. The MCMC chain was run for sufficient length to achieve burn in and effective sample sizes > 200 for all parameters. The maximum clade credibility tree was obtained as described above. We then pruned taxa not included in venom analyses from the tree while retaining the original node ages, and used this tree for all subsequent analyses.

**Venom gland transcriptomics**

Dissected venom glands were homogenized using a pestle and mortar, before RNA was extracted using the TRIzol Plus RNA Purification kit. RNA samples were then DNAse treated and mRNA selected for using the Dynabeads mRNA DIRECT purification kit protocol (Life Technologies). The RNA-Seq libraries were generated from 50 ng of each enriched sample using the TruSeq Stranded mRNA HT Sample Prep Kit (Illumina). As per the Illumina cBot User guide protocol, samples were then denatured and loaded at a concentration of 10 pM, and sequencing was carried out on multiple lanes (six samples multiplexed per lane) of an Illumina MiSeq with 2 × 250 bp paired-end sequencing (Centre for Genomic Research, University of Liverpool). The resulting reads were quality processed using CGR's standard protocols (*11*, *16*, *39*), and then assembled into contigs using the transcriptome assembly program VTBbuilder (*40*). The following parameters were used for assembly: minimum transcript length 150 bp, minimum read length 150 bp, minimum isoform similarity 96%. As part of this process, the VTBuilder algorithm performs normalized read mapping approach on the assembled contigs to generate relative transcript expression data, which is expressed for each contig as a percentage of the total transcriptome expression (*40*). The resulting assembled contigs were then converted into six frame translations, to provide amino acid databases for later protein spectrum matching. Assembled DNA contigs were also annotated with BLAST2GO Pro v3 (*41*) using a BLASTx algorithm and a threshold of 1e-3 against the NCBI non-redundant protein database (GenBank release 219). Manual curation of BLAST2GO annotations were undertaken using the BLASTx annotations on all contigs with expression levels of 0.1% of the transcriptome or higher. Based on these annotations, contigs were classified into toxins (those showing homology with known snake venom toxins previously identified in the literature), non-toxins or ‘unknown’ (if no BLAST annotation or only hypothetical matches were detected). Toxins were further classified by toxin family and the contigs of all toxin families with >1% expression in at least one cobra species were subjected to downstream analysis in MEGA v7 (*42*). As a quality control step, contigs were removed from these datasets if any of the following conditions were not met: i) sequence length was less than 100 bp, ii) 50% or more of the sequence did not match the target toxin in a BLASTx search, iii) the first 50% of the sequence was interrupted by a stop codon, indicating the presence of a pseudogene or misassembly, or iv) the sequence was made up of two exonic regions interspersed by an intron, indicating genomic DNA contamination. For some sequences, it was apparent that ‘underclustering’ had occurred during the assembly process; a common by product of assembling isoform rich gene data (*40*), but more desirable than the alternative scenario of frequently producing chimeric contigs. To resolve this issue, contigs were merged if a ≥50 bp overlap resulted in ≥98% similarity. Sequences were then trimmed to the open reading frame and aligned using the MUSCLE algorithm (*43*) in MEGA’s amino acid space, before the alignments were visually inspected for errors, and edited manually if required.

**Top-down venom proteomics**

Denaturing top-down proteomic experiments were performed as described in previous papers (*44*, *45*). In short, venom samples were dissolved in ultrapure water to a final concentration of 10 mg/mL, and centrifuged at 12,000 x g for 5 min. For reduction of disulfide bonds, 10 µL of dissolved venom was mixed with 10 µL of 0.5 M tris(2-carboxyethyl) phosphine (TCEP), and 30 µL of 0.1 M citrate buffer (pH 3). After 30 min incubation at 65 °C, samples were mixed with 50 µL of acetonitrile/formic acid/$H_2O$ (10:1:89, v/v/v) and centrifuged at 12,000 x g for 5 min. After centrifugation, 5 μL of supernatant of reduced samples were injected for LC-MS/MS analyses. LC-MS/MS experiments of two technical replicates were carried out on a Vanquish ultra-high-performance liquid chromatography (UHPLC) system coupled to a Q-Exactive hybrid quadrupole orbital ion trap (Thermo Fisher Scientific, Bremen, Germany). LC separation was performed on a Supelco Discovery Biowide C18 column (300Å pore size, 2 x 150 mm column size, 3 μm particle size) at a temperature of 30 °C. A flow rate of 0.5 mL/min was used and the samples were eluted with a gradient of water with 0.1% formic acid (FA) (solution A) and 0.1% FA in acetonitrile (ACN) (solution B). Gradient elution started with an isocratic phase with 5% B for 0.5 min, followed by a linear increase to 40% B for 50 min, and 40-70% B for 60 min. Finally, the column was washed with 70% B for 5 min and re-equilibrated at 5% B for 5 min. ESI settings of the mass spectrometer were adjusted to 53 L/min sheath gas, 18 L/min auxiliary gas, spray voltage 3.5 kV, capillary voltage 63 V, S lens RF level 90 V, and capillary temperature 350 °C. MS/MS spectra were obtained in data dependent acquisition (DDA) mode. Mass spectra were acquired with 1 micro scan and 1000 ms maximal C-trap fill time. AGC targets were set to 1E6 for MS1 full scans and to 3E5 for MS/MS scans. Both, the survey scan and data dependent MS/MS scans were performed with a mass resolution (R) of 140,000 (at m/z 200). For MS/MS, the three most abundant ions of the survey scan with known charge were selected for Higher-energy C-trap dissociation (HCD) at the apex of a peak within 2 to 15 s from their first occurrence. Normalized collision energy was stepwise increased from 25% to 30% to 35%. The default charge state was set to z = 6, and the activation time to 30 ms. The mass window for precursor ion selection was set to 3 m/z. A window of 3 m/z was set for dynamic exclusion within 30 s. Ion species with unassigned charge states as well as isotope peaks were excluded from MS/MS experiments.

For data analysis, LC-MS/MS .raw files were converted to .mzXML file format using MSconvert of the ProteoWizard package (version 3.065.85). Extracted ion chromatograms (XICs) of intact proteins were generated of deconvoluted of multiple charged spectra with XTRACT of the Xcalibur Qual Browser version 2.2 (Thermo, Bremen, Germany). XICs of mono-isotopic deconvoluted LC-MS runs were performed with MZmine 2 (version 2.2). A 1.0E4 signal intensity threshold was used. The mass alignment for the creation of XICs was performed with a minimum peak width of 30 s, and 3.0E4 peak height. Mass tolerance was set to 10 ppm. For chromatographic deconvolution, the baseline cutoff algorithm with 1.0E4 signal threshold was used. Maximum peak width was set to 10 min. Feature alignment was performed with 10 ppm mass accuracy and 0.5 min retention time tolerance. For protein spectrum matching, multiple charged MS/MS spectra were then deconvoluted using MS-Deconv (version 0.8.0.7370). The maximum charge was set to 30, maximum mass was set to 50,000, signal-to-noise threshold was set to 2, and m/z tolerance was set to 10 ppm. Protein spectrum matching was performed using TopPIC (http://proteomics.informatics.iupui.edu/software/toppic/) (*46*) (version 1.1.0) against the corresponding venom gland transcriptomic derived protein sequence database as well as all protein sequences from members of the genus *Naja* obtained from the NCBI protein database (Nov 2017), containing a total of 1,764 sequences. The search database also contained 116 protein sequences of known typical contaminants from the common Repository of Adventitious Proteins, (cRAP). TopPIC mass error tolerance was set to 10 ppm and a 1% false discovery rate (FDR, target-decoy) cut-off was used. The maximal allowed unexpected PTMs were set to one. Pairing of MS/MS derived protein ID from TopPic with MS1 XICs was performed by mass matching with an in-house R script available as a jupyter notebook at https://github.com/DorresteinLaboratory/match_tables_by_exact_mass. For relative quantification, area under the curve of XICs was normalized to total ion current (TIC) (see Data S1 and Fig. S2). Amino acid sequences of proteins were converted to FASTA format and imported into MEGA v7 (*42*) for alignment. Proteins were merged if found to be identical using MEGA's pairwise distance measurement.

**Data analyses**

Ancestral states for spitting (vs non-spitting) were estimated over the species tree using the rerooting method function of the R package "phytools" (*47*). After Log-likelihood (Log-lik) comparison of the all rates different (ARD), equal rates (ER) and symmetric rates (SYM) models, both ER and SYM (which had equal Log-lik values) were used to plot these discrete traits over the tree, with both resulting in visually-identical plots. Ancestral states for all continuous characters were also estimated via maximum likelihood on a pruned version of this species tree, using the contmap function of the phytools package under default settings. To test for statistical support for differences between spitting and non-spitting cobras and the influences of geography at the proteomic and functional levels, we used the phylogenetic Generalized Least Squares (PGLS) approach using the pgls() function of the "caper" (*48*) package in RStudio (*49*), with the formula set as ([toxin characteristic, e.g. $PLA_2$ abundance] ~ grouping [spitter or non-spitter]) and lambda set to "Maximum Likelihood". Analyses were performed on the species tree pruned to those species whose venom was analysed in this study. To test for the clustering of spitting and non-spitting lineages based on the amino acid sequences of toxin proteins, we used Principal Coordinate Analysis (PCoA). To do this, a pairwise distance matrix was firstly conducted on the toxin amino acid sequences using the JTT matrix-based model (*50*) in MEGA v.7. The rate variation among sites was modelled with a gamma distribution (shape parameter = 0.8657) for

$PLA_2$s and uniform rates for CTXs. A Euclidean dissimilarity matrix was generated from each of these matrices using the Daisy() function from the R package "Cluster" (*51*) and was followed by applying classical multidimensional scaling to the matrix using the cmdscale() function in R Studio.

**Phylogenetic analyses**

Phylogenetic analysis of three-finger toxin (3FTX) gene family was conducted using Bayesian inference on the aligned sequences derived from the venom gland transcriptomes. Construction of an initial 3FTX tree also used representative sequences from *Aspidelaps scutatus intermedius* (*16*), *Bungarus flaviceps* (*52*), *Bungarus multicinctus* (52), *Dendroaspis* spp. (*11*), *Micrurus fulvius* (*55*) and *Ophiophagus hannah* (53, 54) for phylogenetic context, and a non-toxic *Python regius* 3FTX sequence (*57*) was used as the outgroup for rooting the tree. The final DNA alignment consisted of 56,781 bp with 285 sequences from 28 taxa. The selected model for nucleotide sequence evolution, "GTR+G", was determined by jModelTest v2.1.6 based on the Akaike Information Criterion (*58*, *59*). Subsequently, Bayesian inference phylogenetic analysis was performed using MrBayes v3.2.6 (*60*) on the CIPRES Science Gateway (*61*). The analysis used four simultaneous runs with four chains (three hot, one cold) for $10\times10^6$ generations, sampling every 500th cycle from the chain and using default settings for priors. Burn-in was set at 25%; trees generated prior to this point were discarded, and a consensus tree constructed from the remaining 75%. From the resulting initial 3FTX tree, a strongly supported CTX clade was identified via BLASTx searches on individual sequences and a separate pared-down CTX-specific analysis was also performed to explore the evolutionary history of CTXs sequences in elapids. In addition to the CTXs identified in the original analysis, this second dataset included CTX sequences from *Ophiophagus hannah*

(53, 54) and used the non-CTX 3FTX from *Bungarus flaviceps* (GU190795.1) (*52*) as the outgroup for rooting. The resulting DNA alignment consisted of 18,098 bp with 80 sequences from 17 taxa, phylogenetic analyses were performed as described above, except using the selected "TVM+G" model of nucleotide substitution, selected using the same methods described above in jModelTest. The two sequence alignments are displayed in Data S2 and S3, and the resulting phylogeny of the CTX-specific analysis is displayed in Fig. S3.

**Venom cytotoxicity via hen's egg test-chorioallantoic membrane (HET-CAM) assay**

Lyophilized venom (Table S10) was dissolved in sterile, deionized water to make a stock solution. The protein concentration was determined using a NanoDrop 1000 UV/Vis Spectrophotometer (Thermo Fisher Scientific, Bleiswijk, the Netherlands) at an absorbance of 280 nm. The stock solution was snap-frozen using liquid nitrogen and stored at −80°C till further use. Just before the experiment the venom was prepared in Hanks' balanced salt solution (HBSS: Cat. H9394, Sigma Aldrich, Zwijndrecht, the Netherlands) to a final total protein concentration of 1 mg/mL. Unincubated hen's eggs were obtained from a commercial supplier. The eggs were incubated in a horizontal position at 38°C on stationary shelves in a humidified incubator for 10 days. Following incubation, eggs were candled and placed vertically with the blunt end upwards. A hole in the eggshell overlying the air sac was made with No. 3 watchmakers' forceps. The shell membrane was moistened with 200 µL HBSS, and then punctured and carefully peeled away using No. 5 watchmakers' forceps to reveal the chorioallantoic membrane (CAM) below. Excess HBSS was pipetted away. Venom or control solution (100 µL) was then pipetted onto the CAM and a timer started. Any hyperaemia, haemorrhage and/or coagulation was documented over a 5 min period. Photographs were taken at 15 s intervals for semi-quantitative analysis. Each venom and control sample was tested in triplicate (using three different eggs) in a single-blinded and randomized manner to avoid bias. The positive controls were 1% sodium dodecyl sulphate (Carl Roth, Karlsruhe, Germany) and 1 mg/mL of capsaicin (Sigma Aldrich, Zwijndrecht, the Netherlands), both known eye-irritants; and 10 mg/mL of histamine (Sigma Aldrich, Zwijndrecht, the Netherlands), a potent vasodilator. The negative control was HBSS. A semi-quantitative analysis was performed using the resulting photographs, where the severity of any hyperaemia, haemorrhage and/ or coagulation was manually scored at 0.5, 2.0 and 5.0 min using the method developed by Luepke (*62*) (Tables S3, S4). The average score of three replicates was taken and the cumulative score was matched with the corresponding 'irritation potential' previously developed (*62*) (Table S4).

**Calcium imaging on mouse sensory neurons**

Trigeminal ganglia (TG) or dorsal root ganglia (DRG) from 4-6 week old male C57BL/6 mice were dissociated and plated in DMEM (Gibco, MD, USA) containing 10% fetal bovine serum (FBS) (Assaymatrix, VIC, Australia) and penicillin/streptomycin (Gibco, MD, USA) on a 96-well poly-D-lysine-coated culture plate (Corning, ME, USA) and maintained overnight. Cells were loaded with Fluo-4 AM calcium indicator, according to the manufacturer's instructions (ThermoFisher Scientific, MA, USA). After loading (1 h), the dye-containing solution was replaced with assay solution (1x Hanks' balanced salt solution, 20 mM HEPES). Images were acquired at 20x objective at 1 frame per second (excitation 485 nm, emission 521 nm). Fluorescence corresponding to $[Ca^{2+}]_i$ of 100–150 cells per experiment was monitored in parallel using an Nikon Ti-E Deconvolution inverted microscope, equipped with a Lumencor Spectra LED Lightsource. Baseline fluorescence was monitored for 30 s. At 30 s, assay solution was replaced with venom (100 ng/μL in assay solution) or venom fractions (estimated equivalent of amount for 100 ng/μL whole venom, in assay solution) and monitored for an additional 2 min. Experiments involving the use of mouse tissue were approved by The University of Queensland animal ethics committee.

**FLIPR assays of F11 cells**

F11 (Neuroblastoma × dorsal root ganglion (DRG) neuron hybrid) cells were cultured as previously described (*63*). Briefly, cells were maintained on Ham's F12 media supplemented with 10 % fetal bovine serum, 100 μM hypoxanthine, 0.4 μM aminopterin, and 16 μM thymidine (Hybri-MaxTM, Sigma Aldrich). A 384-well imaging plate (Corning, Lowell, MA, USA) was seeded 48 h prior to calcium imaging resulting in 90 – 95% confluence at imaging. Cells were incubated for 30 min with Calcium 4 assay component A, according to manufacturer's instructions (Molecular Devices, Sunnyvale, CA) in physiological salt solution (PSS; composition in mM: 140 NaCl, 11.5 D-glucose, 5.9 KCl, 1.4 $MgCl_2$, 1.2 $NaH_2PO_4$, 5 $NaHCO_3$, 1.8 $CaCl_2$, 10 HEPES) at 37 °C. $Ca^{2+}$ responses were measured using a FLIPRTETRA fluorescent plate reader equipped with a CCD camera (Ex: 470-490 nm, Em: 515-575 nM) (Molecular Devices, Sunnyvale, CA). Signals were read every second for 10 s before, and 300 s after the addition of crude venoms (final concentration range: 1.3 – 333 ng/μL) in PSS supplemented with 0.1% bovine serum albumin. All data represent the mean ± standard error of the mean (SEM) of a representative assay in triplicate unless otherwise stated. The mean fluorescence changes during venom addition was compared to a control solution addition, (0.1% BSA in PSS). The resulting maximum-minimum florescence in the 300 s period after venom was added was recorded as the response. A four-parameter Hill equation (Variable slope, two-site) was fitted using Graphpad Prism8.

**Venom fractionation and bottom-up sequencing**

To identify the venom constituents responsible for sensory neuron activation, 500 µg of venoms from *N. siamensis*, *N. nigricollis* and *H. haemachatus* were each separated on a Phenomenex Gemini NX-C18 column (250 x 4.6 mm, 3 µm particle size, 110 Å pore size) using a gradient of 15–45% solvent B (90% ACN, 0.05% TFA) over 30 min at a flow rate of 1 mL $min^{-1}$. Fractions were collected on the basis of absorbance at 214 nm. Fractions were dried by vacuum concentration and each resuspended in 50 µL pure water from which 1 µL aliquots were used for calcium imaging experiments (diluted in 100 µL assay solution). Bottom-up proteomics was used to confirm the identity of the major component/s of each fraction. 20 µL of each resuspended fraction was dried by vacuum centrifugation. Each sample was reduced and alkylated with reagents in gas phase according to the protocol described by Hale et al. (*64*). Reduction/alkylation reagent (50% 0.1 M ammonium carbonate, 48.75% ACN, 0.25% triethylphosphine, 1% 2-iodoethanol (by volume)) was placed in the cap of each inverted tube and incubated at 37 °C for 60 min. Reduced and alkylated fractions were then digested by incubating with 20 ng/µL trypsin overnight at 37 °C according to the manufacturer's instructions (Sigma-Aldrich, MO, USA). Trypsin digestion was quenched by addition of FA to a final concentration of 0.5%. Each sample was then separated on a Shimadzu (Japan) Nexera uHPLC with an Agilent Zorbax stable-bond C18 column (2.1 mm × 100 mm, 1.8 µm particle size, 300 Å pore size), using a flow rate of 180 µL/min and a gradient of 1–40% solvent B (90 % ACN, 0.1% FA) in 0.1% FA over 30 min and analyzed on an AB Sciex 5600 TripleTOF mass spectrometer. MS survey scans were acquired at 300–1800 m/z over 250 ms, and the 20 most intense ions with a charge of +2 to +5 and an intensity of at least 120 counts were selected for MS/MS. The unit mass precursor ion inclusion window mass ± 0.7 Da, and isotopes within ±2 Da were excluded from MS/MS, with scans acquired at 80–1400 m/z over 100 ms and optimized for high resolution. Using ProteinPilot version 4.0 (ABSciex), MS/MS spectra were searched against a database of the toxin sequences (translated open-reading frames) derived from the corresponding venom gland transcriptomes.

**Phospholipase $A_2$ assay**

We characterized the enzymatic $PLA_2$ activity of each venom using a recently described colorimetric assay (*65*). A $PLA_2$ reaction solution was made using 49.5 µM Cresol red dye, 0.875 mM Triton X-1007 M, and 1 mL of 5x Salt Mix (1mM Tris base pH 8.5, 500 mM sodium chloride, 500 mM potassium chloride and 50 mM calcium chloride). The pH of the reaction solution was checked using a pH strip and corrected (if necessary) with 1 M sodium hydroxide. To this solution 168.25 µL of the substrate (L-α-phosphatidylcholine; stock concentration 26 mM, SIGMA Life Sciences) was added. Venom samples were prepared in tenfold dilutions (0.1-100 µg/mL) of Tris buffer (pH 8.5), with 10 µL pipetted into the wells of a Greiner Bio-One clear 384-well microplate at volumes in quadruplicate using a 230V Multidrop 384 (Labsystems), with control wells containing Tris buffer only. The plate was then overlaid with 40 µL of reaction solution in each well and read kinetically on a FLUOstar Omega (BMG LABTECH) spectrophotometer at 572 nm over 42 cycles (~46 s/cycle) at 25 °C. Concentration curves were generated for each venom by plotting the mean control AUC of absorbance at 572 nm, minus the mean area under the curve (AUC) for readings at each venom concentration, using the values from the run period of 0-10 min (Fig. S12). The resulting mean area under the concentration curves (AUCC) were used for comparisons.

**Venom median lethal doses ($LD_{50}$)**

We calculated the lethal dose 50 ($LD_{50}$) of each of the venoms using a previously established murine model of venom lethality (*66*). Groups of five mixed sex CD-1 mice (18-20 g) received intravenous (i.v.) injections, in the caudal vein, of various doses of each venom (Table S5), dissolved in 0.1 mL of 0.12 M NaCl, 0.04 M phosphates, pH 7.2 (PBS). Deaths were recorded at 24 h and the $LD_{50}$, as well as the 95% confidence limits, were estimated by probits. The experimental protocols for these experiments were approved by the Institutional Committee for the Care and Use of Laboratory Animals (CICUA) of the University of Costa Rica (approval number CICUA 82-08).

**CT imaging of snake fangs**

Fangs were scanned in pairs on a Nikon XTH225ST CT scanner at the University of Bristol. All scans were performed on a rotating target with a tube voltage of 70 kV and beam current of 101 μA with a 1 s exposure per projection with a total of 3141 projections in 360° at a resolution of between 3.9 to 4.5 microns. The CT scan data were aligned, reconstructed and cropped to generate individual tiff image stacks for each fang using VG Studio MAX (Volume Graphics GmbH, Heidelberg, Germany). The tiff stacks were subsequently segmented to generate a 3D model of each fang in Avizo 9 (FEI, Hillsboro, Oregon, USA).

### Supplementary Figures

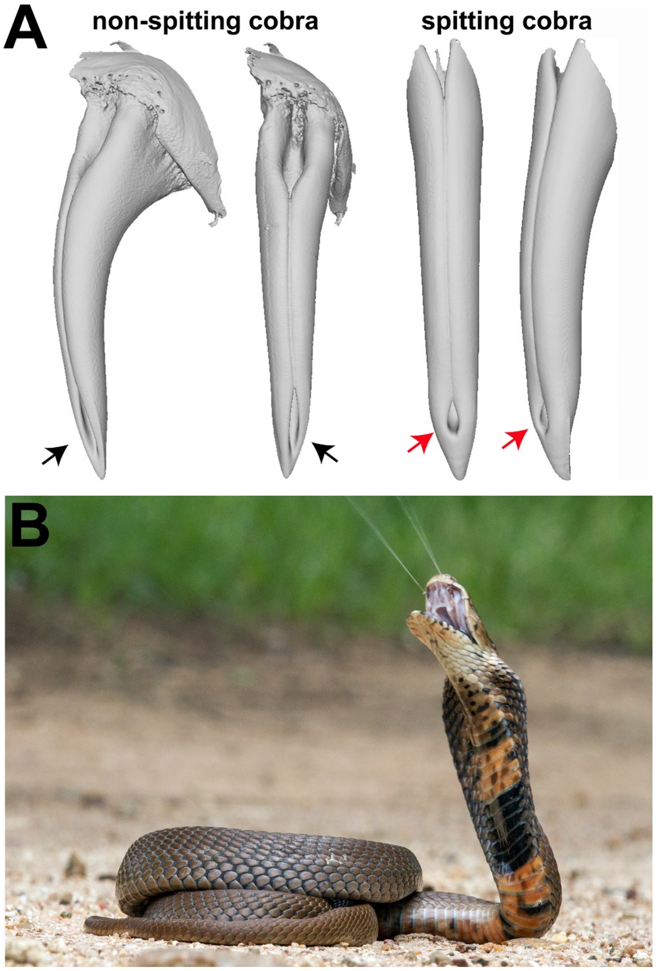


**Fig. S1. The morphological adaptations associated with venom spitting. (**A) MicroCT visualizations of fangs from representative non-spitting (*Naja nivea*) and spitting (*Naja nubiae*) cobra species. Fangs are visualized from the anterior perspective (central images) and rotated 45° (outer images). Black and red arrows highlight the distinct ejection orifices of non-spitting and spitting cobras, respectively. (B) Venom spitting displayed by a *Naja mossambica* from Limpopo Province, South Africa. Photograph by Wolfgang Wüster.

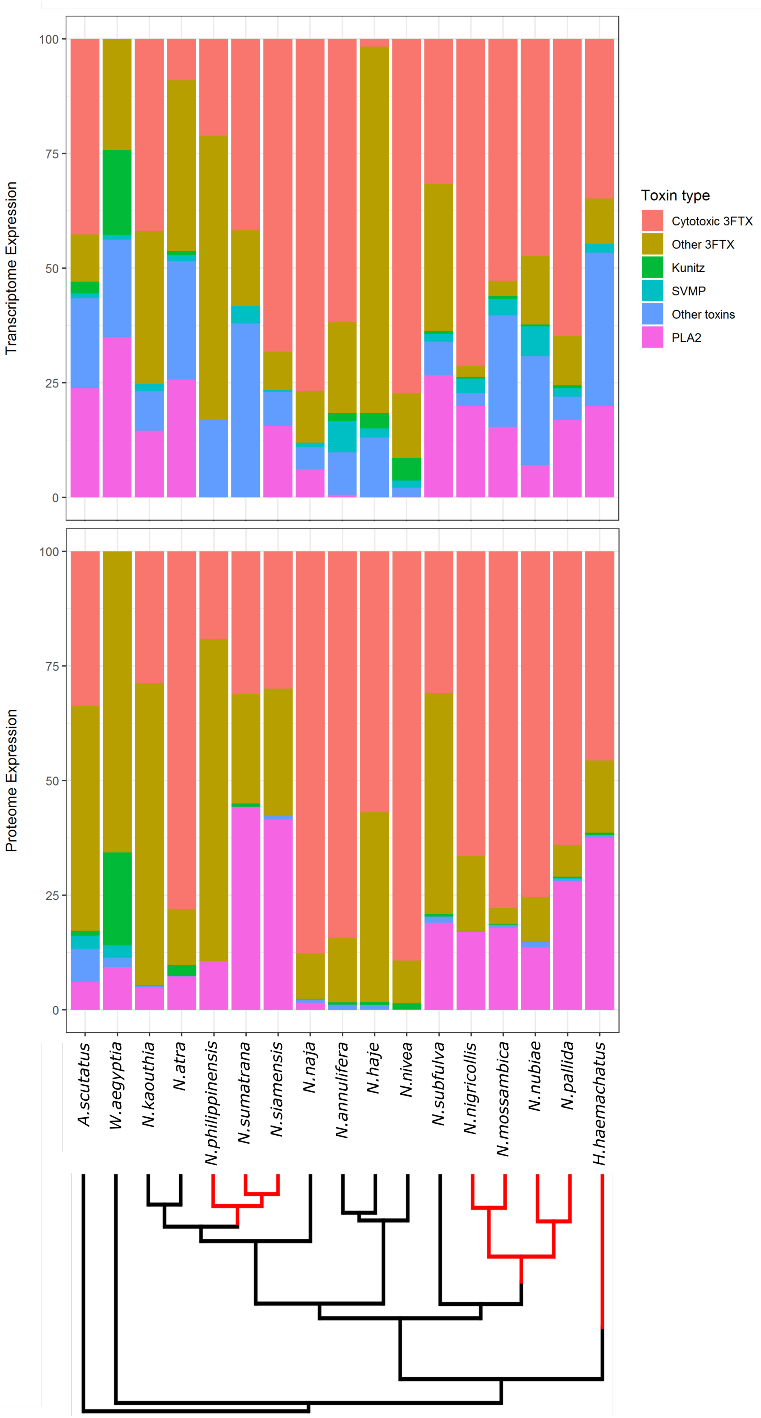


**Fig. S2.** Stacked bar charts representing the relative abundances of toxin families detected in the venom gland transcriptomes (top) and venom proteomes (bottom) of cobras and related species. The transcriptome data presented represents the percentage of all venom toxin-encoding gene expression detected in each venom gland transcriptome based on normalized read mapping to assembled contigs. The proteome data represents the percentage of all venom toxins detected in each venom proteome based on top-down proteomic analyses and quantification via area under the curve of extracted ion chromatograms (XICs) normalized to total ion current (TIC). The bars are plotted over the phylogenetic tree of the species used in this study. Branches of the tree highlighted in red represent spitting lineages. Note that the transcriptomic and proteomic abundances for *A. scutatus* are taken from (*16*), and while the transcriptomic data is directly comparable, the proteomic analyses used a different measure of quantification than that applied for the remainder of the species investigated in this study.

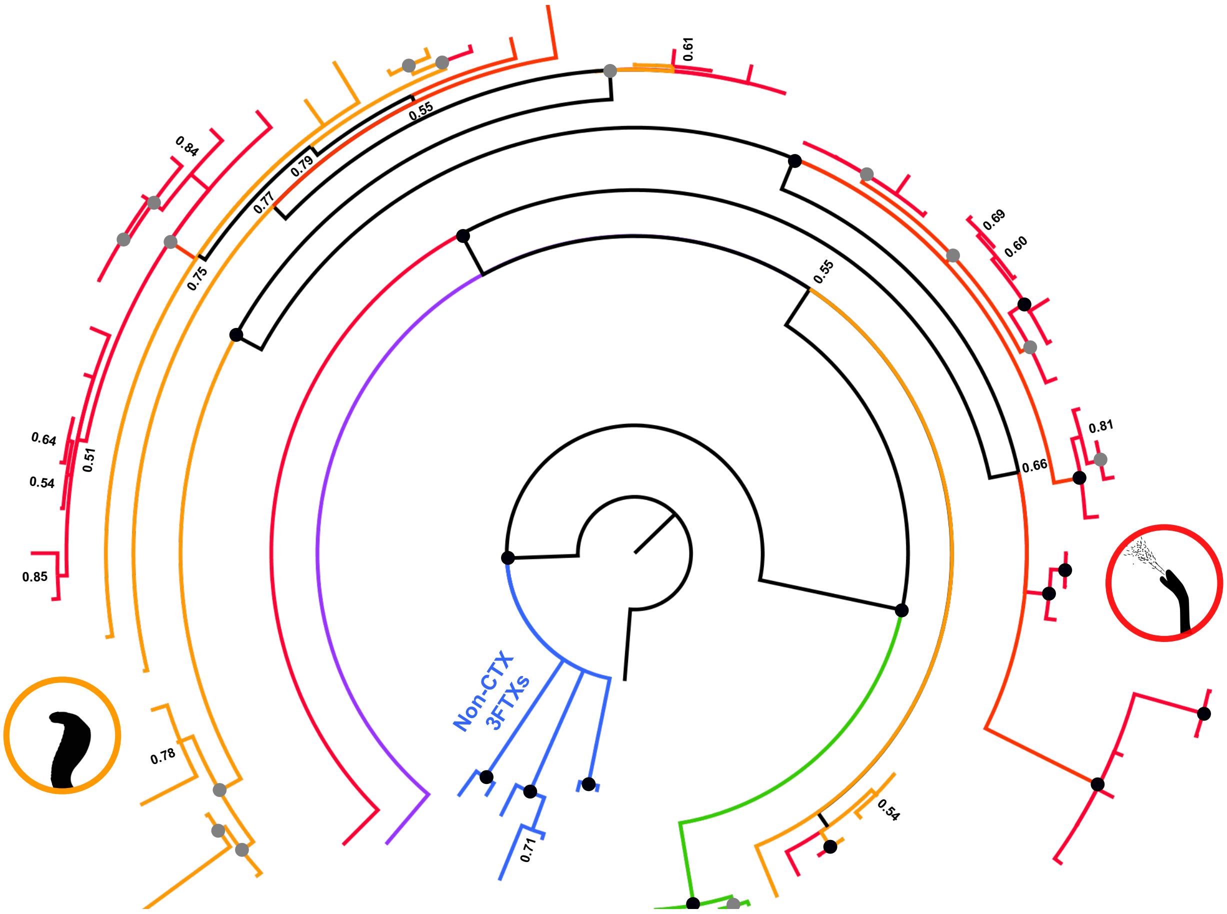


**Fig. S3. Bayesian inference-derived DNA phylogeny of elapid three finger toxin cytotoxin (CTX) sequences demonstrates that the vast majority are from cobra species (genus *Naja*).** Black-filled circles at each node represent Bayesian posterior probabilities of 1 and grey-filled circles represent values between 0.90 and 0.99. Tips labels have been removed to aid visualization. Colored branches represent non-CTX 3FTX outgroups (blue) and CTXs found in: the king cobra *Ophiophagus hannah* (green), *Aspidelaps scutatus* (purple), non-spitting cobras (orange), and spitting cobras (including *H. haemachatus*) (red). No bona fide CTX sequences were detected in the venom gland transcriptome of *Walterinnesia aegyptia*.

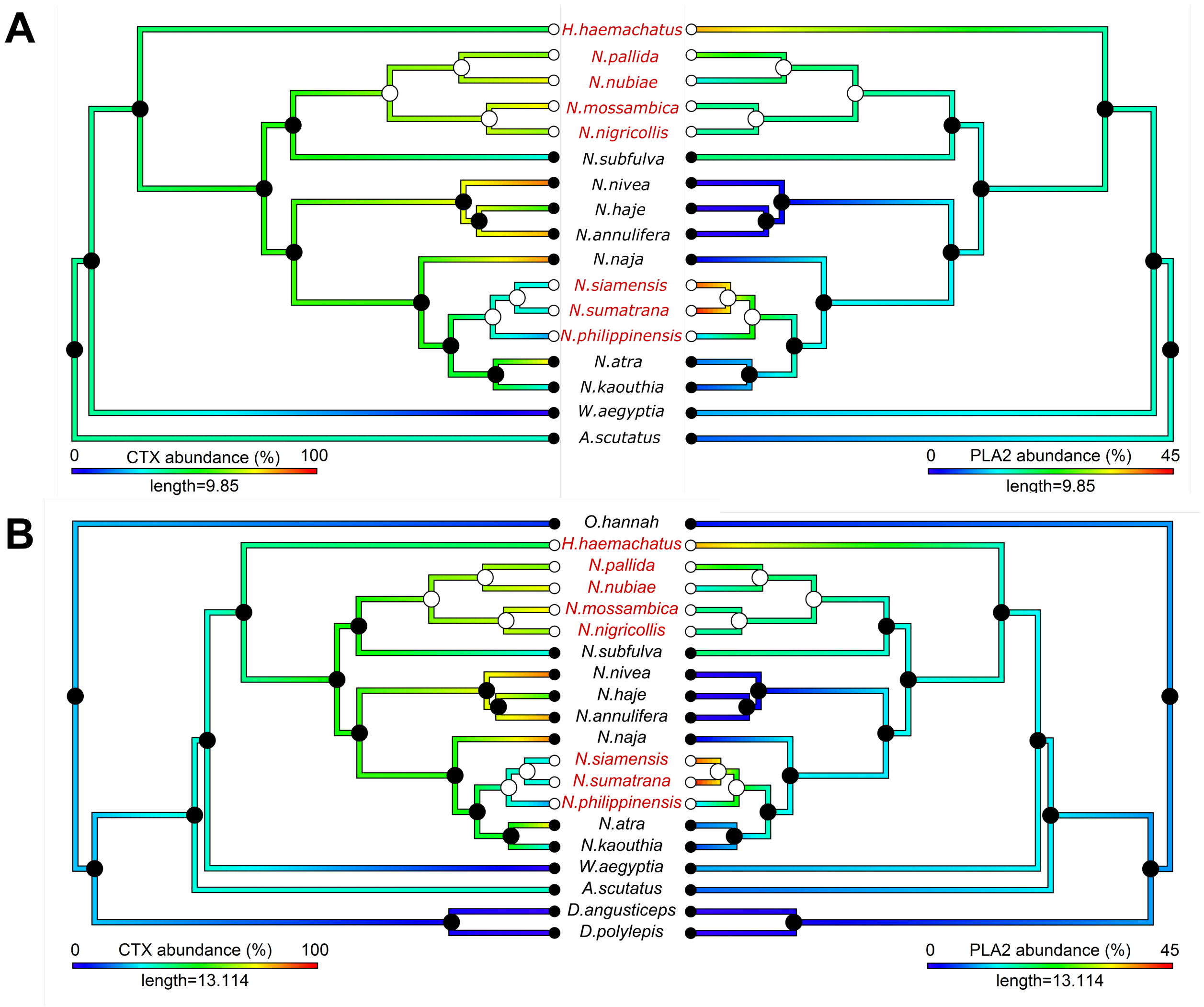


**Fig. S4. Proteomic abundances of phospholipase $A_2$ ($PLA_2$) but not cytotoxic 3FTXs (CTXs) are significantly associated with the emergence of venom spitting.** (A) Ancestral state estimations of the proteomic abundance (percentage of all toxins) of CTXs (left) and $PLA_2$s (right) mapped onto a species tree containing the cobras and representatives of the outgroup genera *Walterinnesia* and *Aspidelaps*. (B) Ancestral state estimations of the proteomic abundance of CTXs (left) and $PLA_2$s (right) mapped onto a species tree containing those taxa displayed in A, as well as the additional more distantly related elapid outgroups *Dendroaspis* (*D. angusticeps* and *D. polylepis*) and *Ophiophagus hannah*. Additional outgroups were incorporated due to the high reconstructed ancestral abundance of CTXs observed in A. For both (A) and (B) filled or empty circles at tips and nodes represent the observed or estimated ancestral state of non-spitting and spitting, respectively. Red tip labels are also used to highlight the spitting lineages. The Phylogenetic Generalized Least Squares (PGLS) statistics for each trait are as follows: CTX abundance with *Aspidelaps* as outgroup (A, left); $t = -0.83$, $df = 15$, $p = 0.42$, and with

*Ophiophagus* as outgroup (B, left); $t = -0.72$, $df = 18$, $p = 0.48$. $PLA_2$ abundance with *Aspidelaps* as outgroup (A, right); $t = 4.27$, $df = 15$, $p = 0.0007$, and with *Ophiophagus* as outgroup (B, right); $t = 5.12$, $df = 18$, $p = 0.00007$. Proteomic abundances for *Aspidelaps scutatus*, *Dendroaspis* spp. and *Ophiophagus hannah* were taken from (*11*, *16*, *67*).

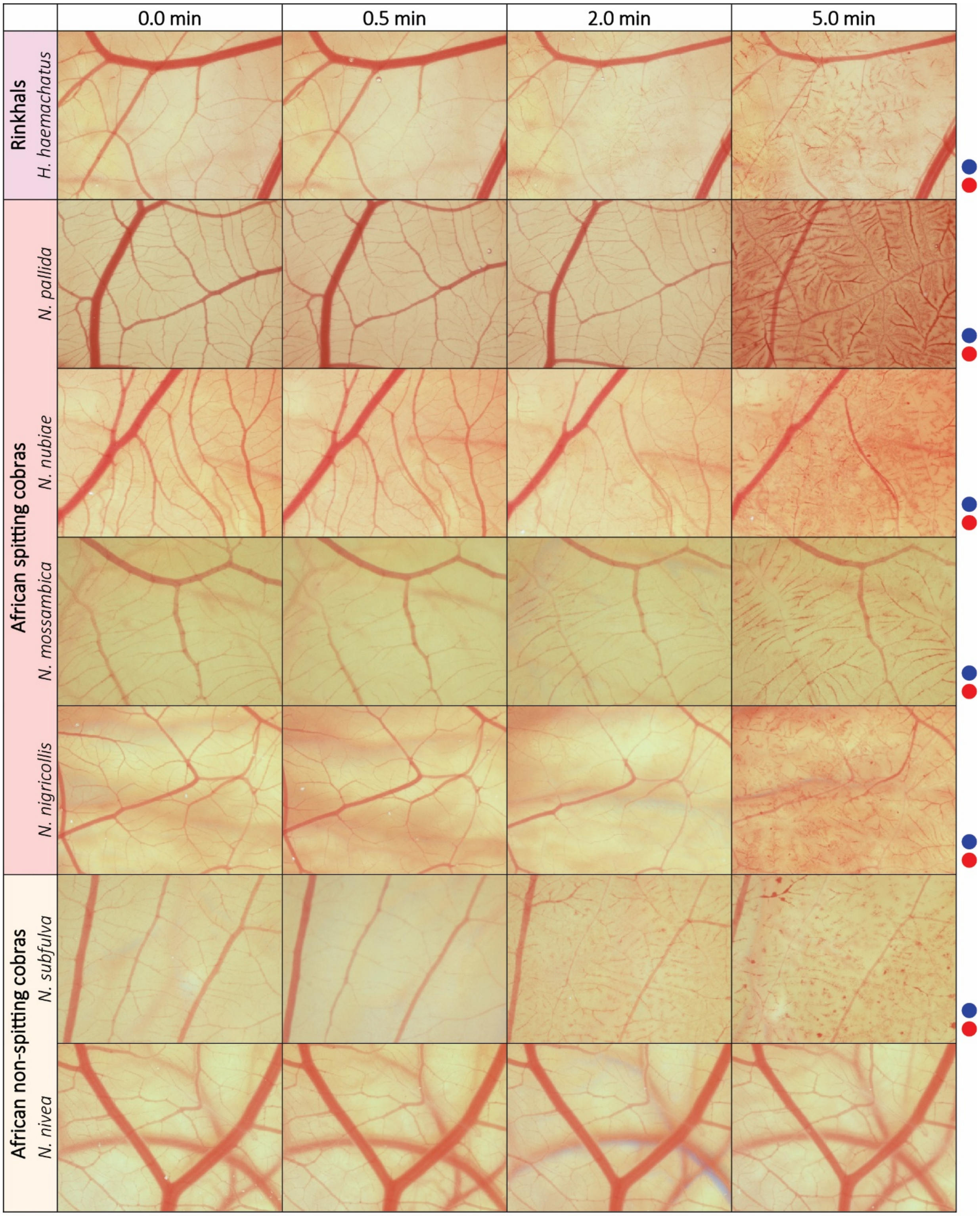
0.0 min
0.5 min
2.0 min
5.0 min
Rinkhals
H. haemachatus
African spitting cobras
N. pallida
N. nubiae
N. mossambica
N. nigricollis
African non-spitting cobras
N. subfulva
N. nivea

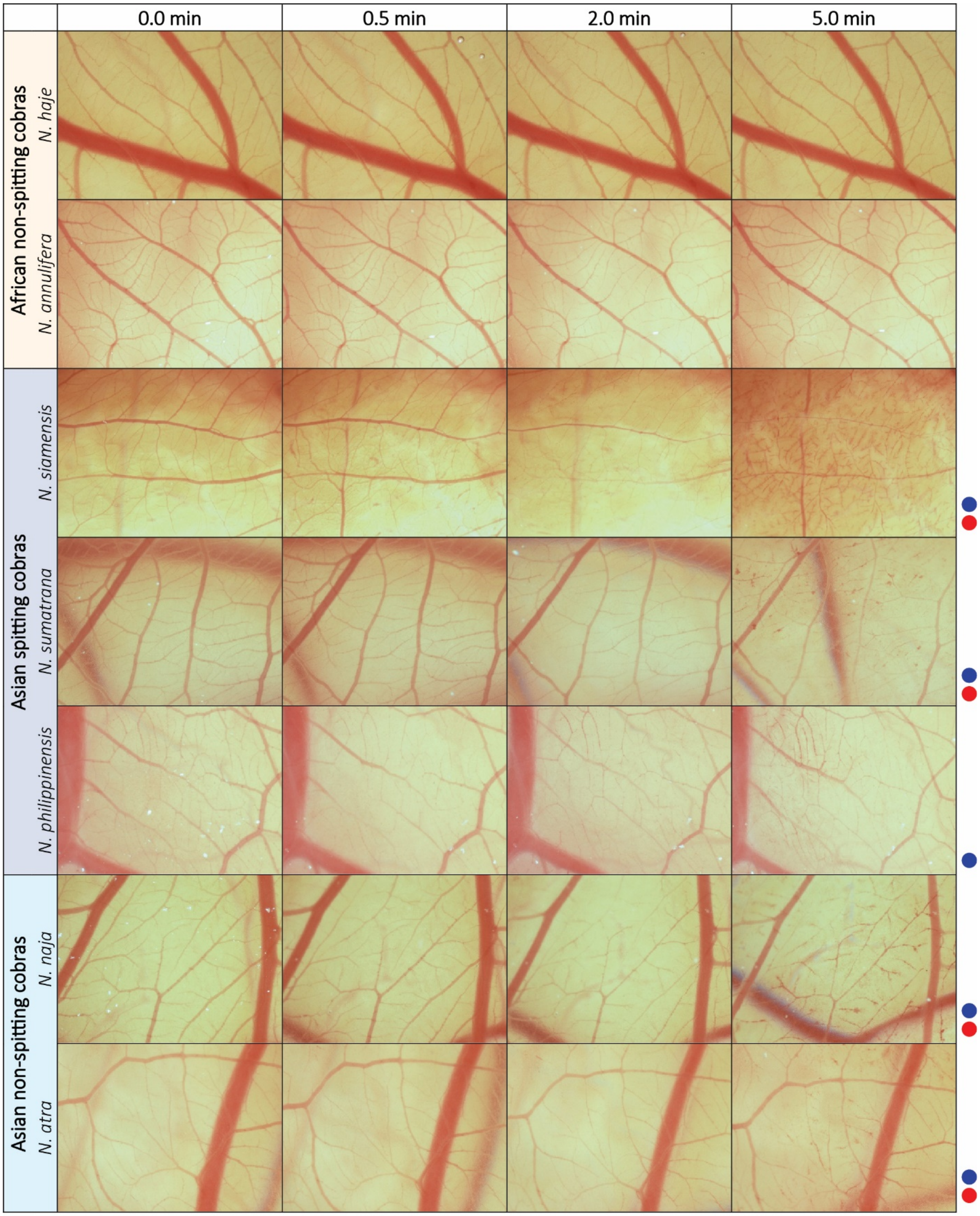

0.0 min
0.5 min
2.0 min
5.0 min
African non-spitting cobras
N. haje
N. annulifera
Asian spitting cobras
N. siamensis
N. sumatrana
N. philippinensis
Asian non-spitting cobras
N. naja
N. atra

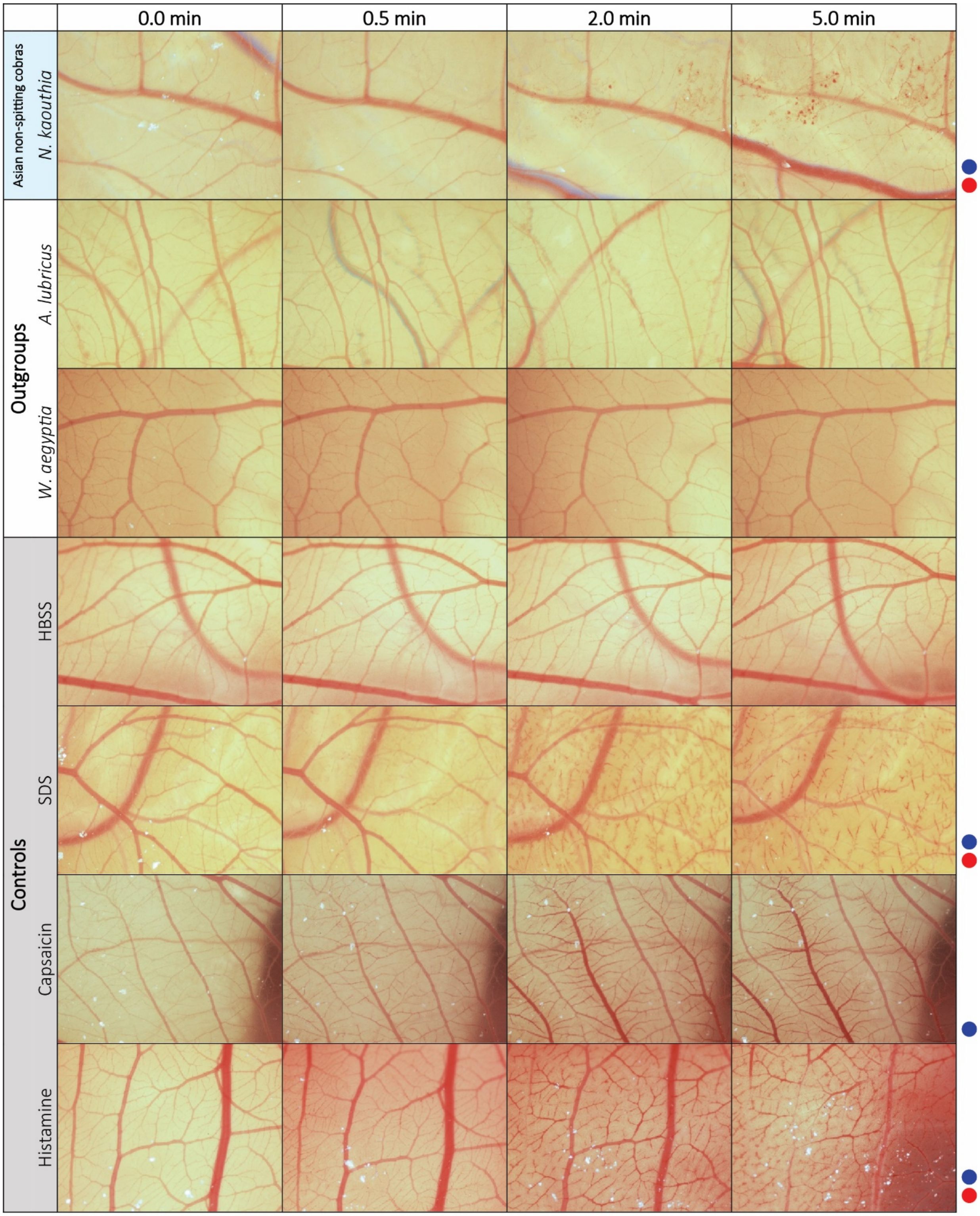


**Fig. S5. The vascular pathologies induced by spitting cobra venoms.** The eye-irritation potential of snake venoms and controls were assessed using the *in vivo* hen's egg test-chorioallantoic membrane (HET-CAM) assay (*62*). The vascular effects observed are indicated on the right of each series if present: hyperaemia (●) and coagulation (●). All series were also assessed for evidence of haemorrhage, but this pathology was not observed in any sample. Key: HBSS, Hanks' balanced salt solution; SDS, sodium dodecyl sulphate.

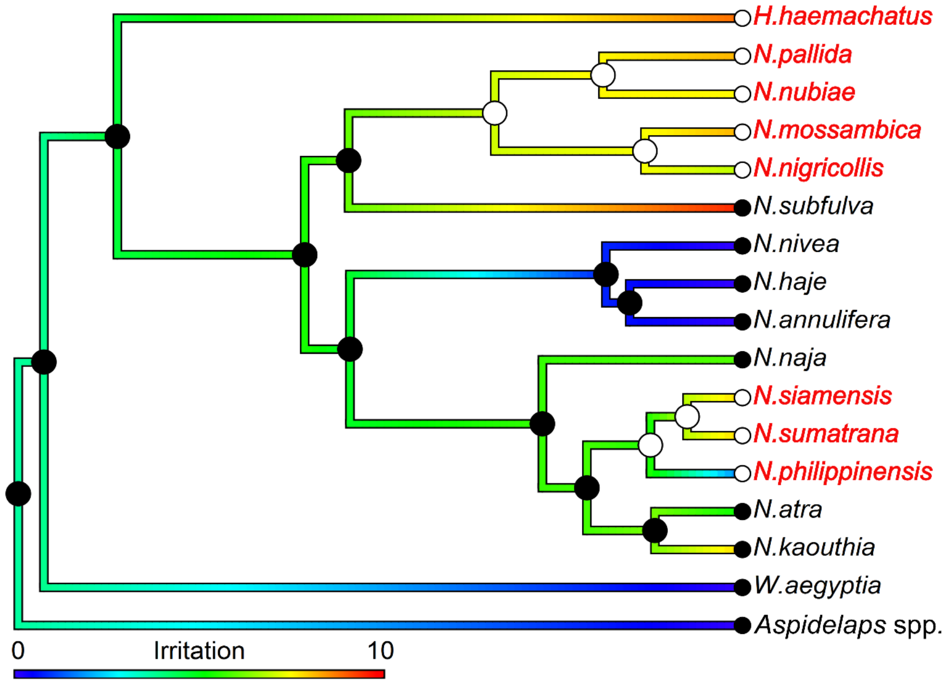


**Fig. S6. Ancestral state estimation of eye-irritation scores, determined in the hen's egg test-chorioallantoic membrane assay, demonstrate no significant association with the emergence of venom spitting.** The Phylogenetic Generalized Least Squares (PGLS) analysis revealed no association between the extent of venom-induced eye irritation and the emergence of spitting ($t = 1.08$, $df = 15$, $p = 0.30$). Colored branches are scaled according to extent of irritation (blue, low abundance; red, high abundance). At each node and tip of the tree, filled or empty circles represent the estimated ancestral state of non-spitting or spitting, respectively, and red tip labels are used to highlight spitting lineages.

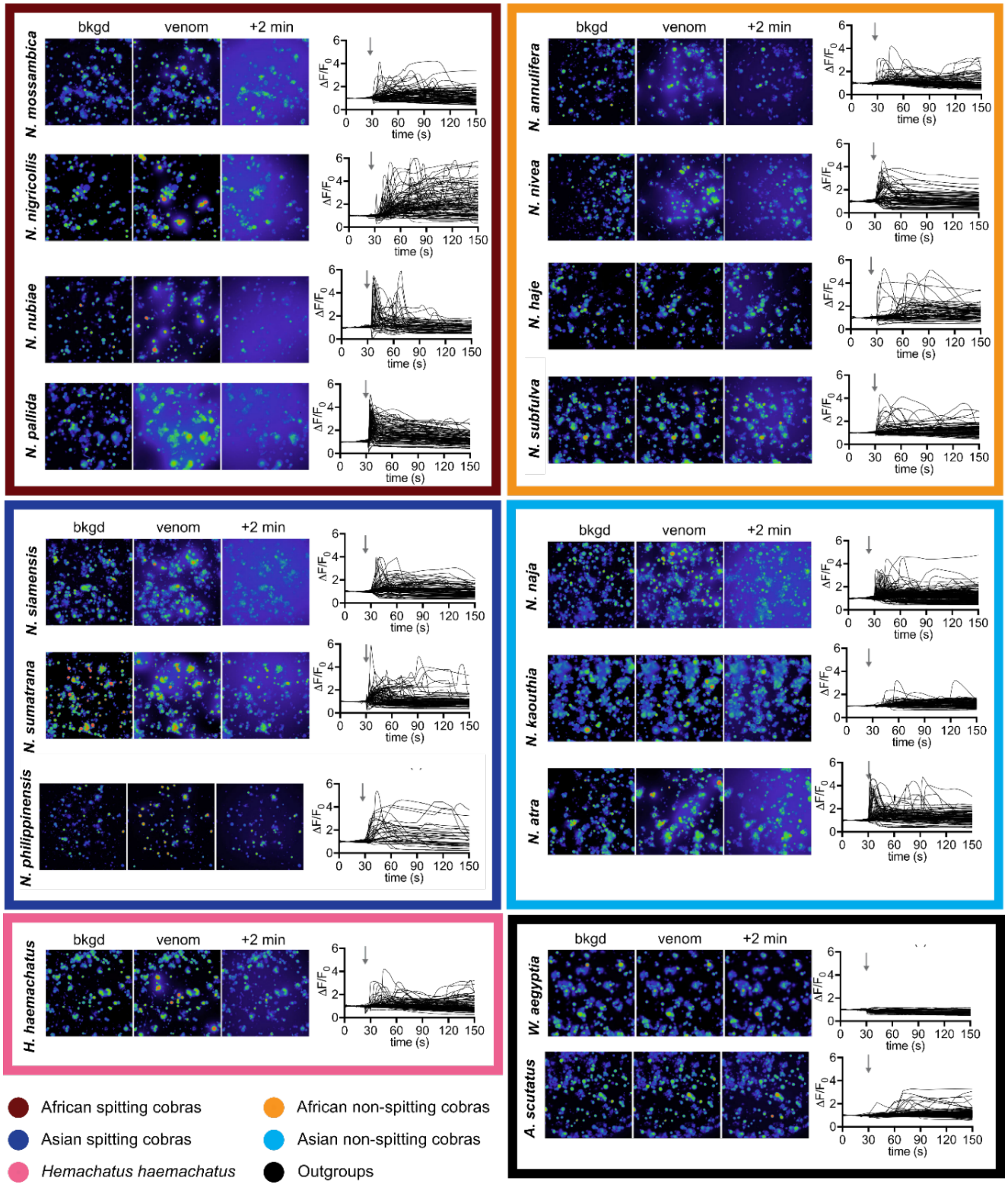


**Fig. S7.** ***Naja* and *Hemachatus* venoms activate sensory neurons *in vitro*.** Application of *Naja* and *Hemachatus* venoms (100 ng/µL) to dissociated mouse trigeminal ganglion cells caused, over the 2 min recording, increases in $[Ca^{2+}]_i$ in both neurons and non-neuronal cells, which was followed by a decrease in $[Ca^{2+}]_i$ reflecting dye leakage from the cells into the media. These data indicated that components of the venom permeate cell membranes allowing a massive influx of $Ca^{2+}$ (and likely other ions) followed by leakage of cellular contents. Such an action on trigeminal ganglion neurons is consistent with the painful effects reported for envenomation.

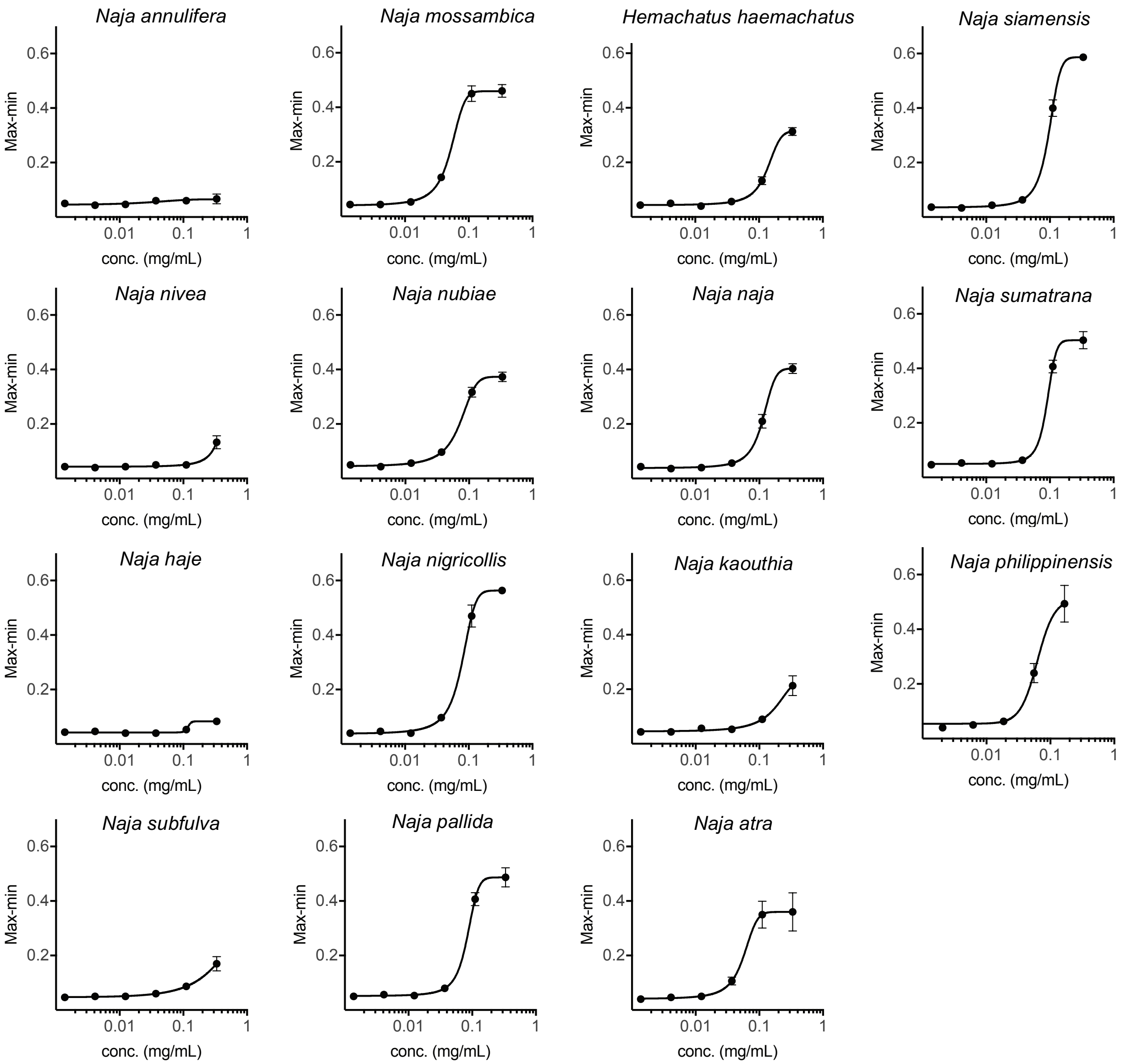


**Fig. S8. Potency of *Naja* and *Hemachatus* venoms on activation of sensory neuron-derived F11 cells.** Values are max-min (n = 3), and error bars represent standard error of the mean (SEM). Curves were fitted using a four-parameter Hill equation (Variable slope) in Graphpad Prism (version 8.02).

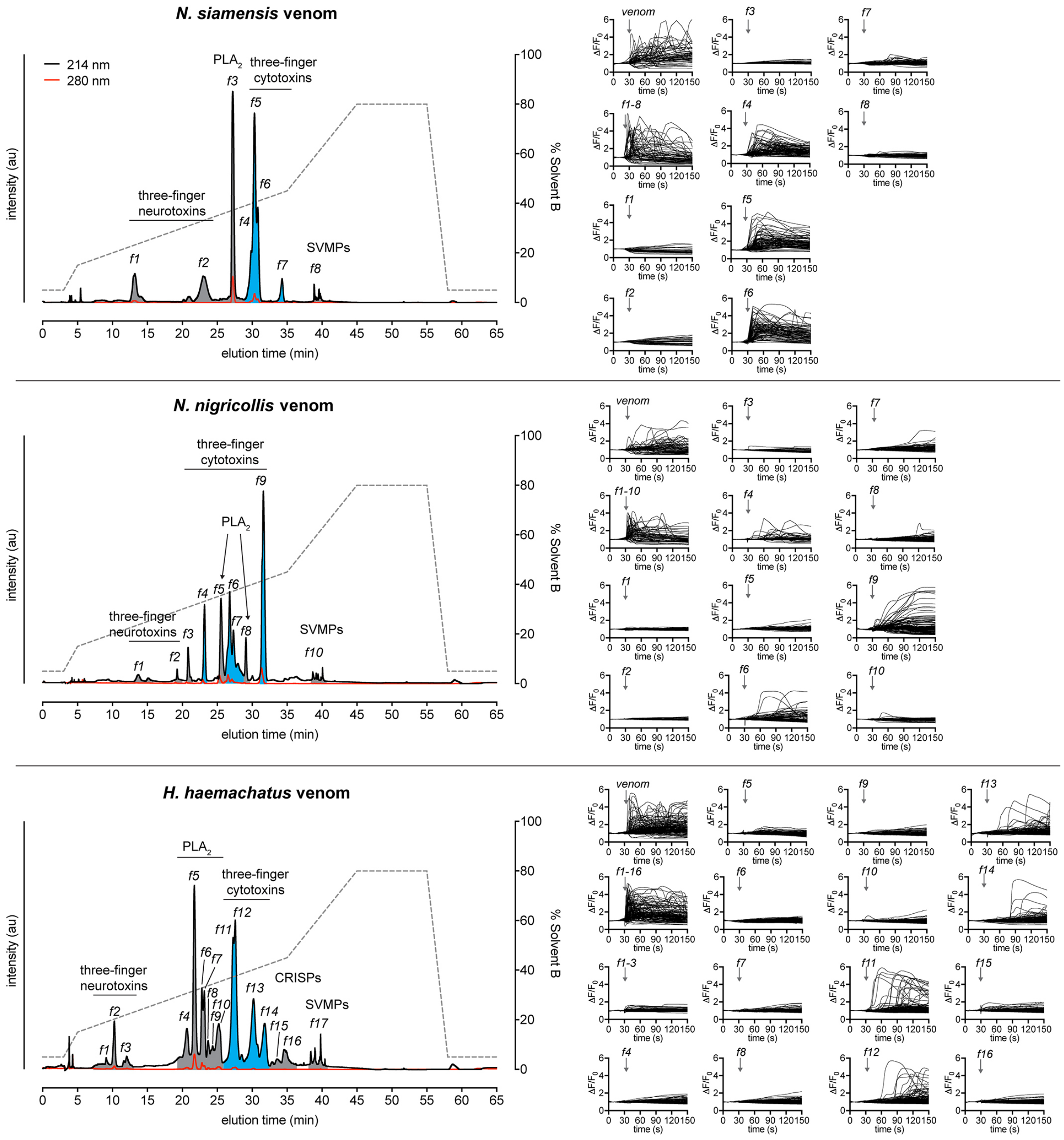


**Fig. S9. Separation of *Naja siamensis, Naja nigricollis* and *Hemachatus haemachatus* venoms and activity of fractions on sensory neurons.** *N. siamensis, N. nigricollis* and *H. haemachatus* venoms (500 µg) were each separated by RP-HPLC using a Phenomenex Gemini NX-C18 column (250 x 4.6 mm, 3 µm particle size, 110 Å pore size; gradient of 15–45% solvent B (90% ACN, 0.05% TFA) over 30 min; flow rate of 1 mL min$^{-1}$). Fractions were collected according to absorbance at 214 nm. Bottom-up proteomics was used to confirm the identity of the major component/s of each fraction. Individual fractions were assessed for activity on sensory neurons (mouse DRG). Active fractions are highlighted in blue, inactive in grey. Each set of traces represents all cells of one experiment. Only the pooled fractions and those individual fractions containing three-finger cytotoxins were able to activate sensory neurons.

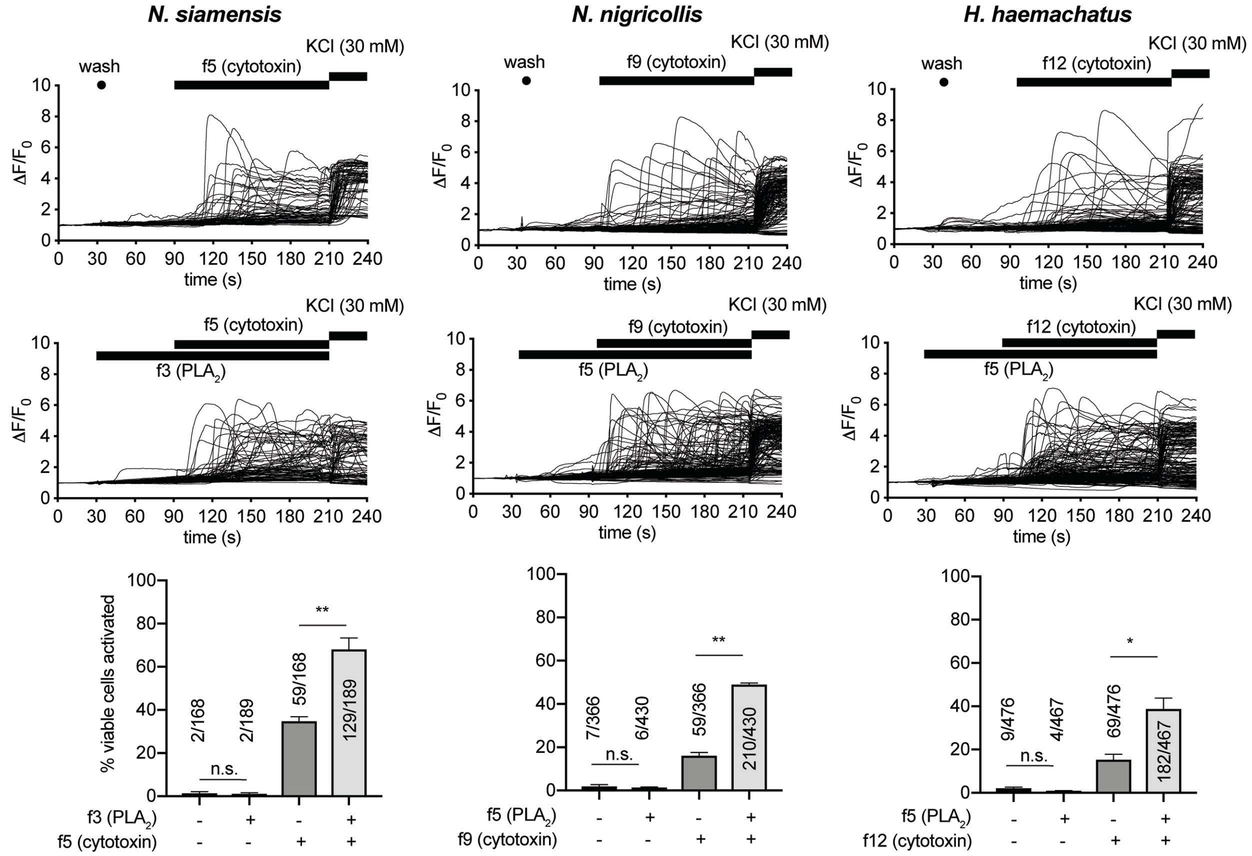


**Fig. S10. Activation of sensory neurons by spitting cobra three-finger cytotoxins is potentiated by $PLA_2$s.** For all venoms tested, there was no significant difference in cell (mouse DRG) activation between buffer/wash and a $PLA_2$ fraction alone, while cytotoxin fraction activation was potentiated in the presence of a corresponding $PLA_2$ fraction. Each set of traces represents all cells of one representative experiment. Data are derived from 2-3 independent experiments. Statistical comparisons were made using unpaired parametric *t*-tests in Graphpad Prism (version 8.02). *, P < 0.05; **, P < 0.01.

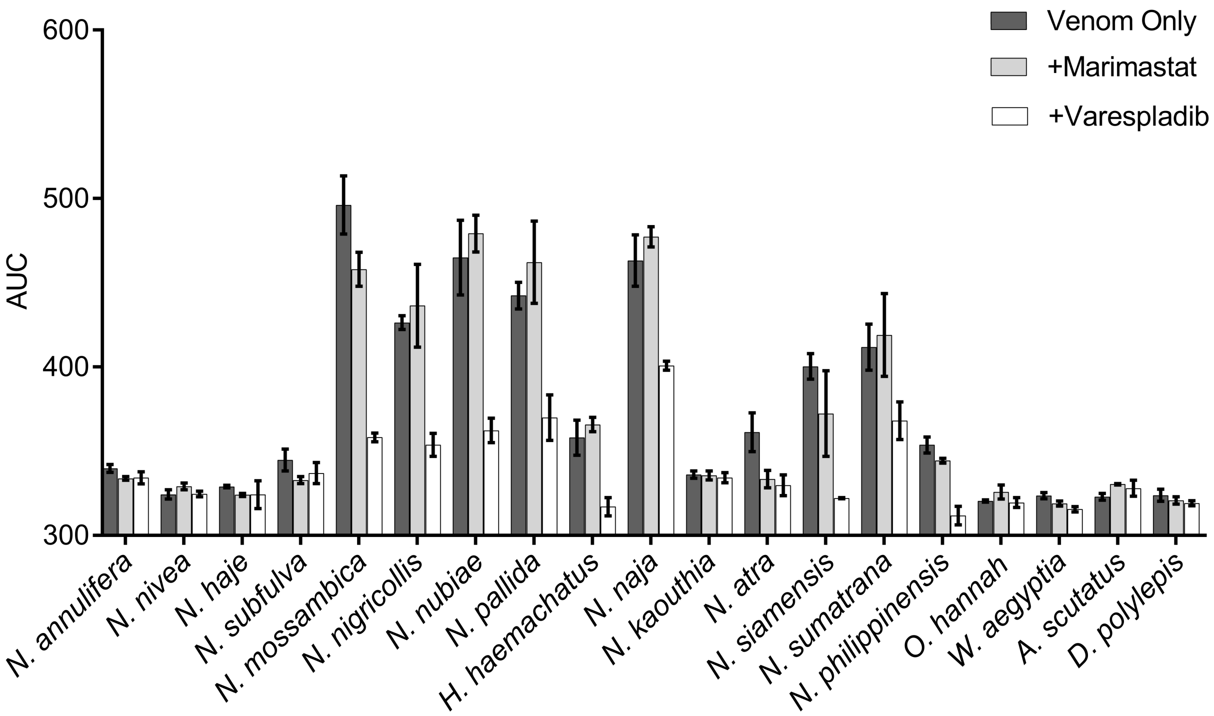


**Fig. S11. The $PLA_2$ inhibitor varespladib reduces the activation of sensory neurons caused by spitting cobra venoms.** 80 µg/mL of venoms (or 160 µg/mL in the case of *H. haemachatus* and *N. philippinensis* venom) were added to F11 cells. Values are area-under-curve (n = 3), and error bars represent standard error of the mean (SEM), with either venom only (negative control), the $PLA_2$ inhibitor varespladib (13 µM) or the metalloprotease inhibitor marimastat (15 µM).

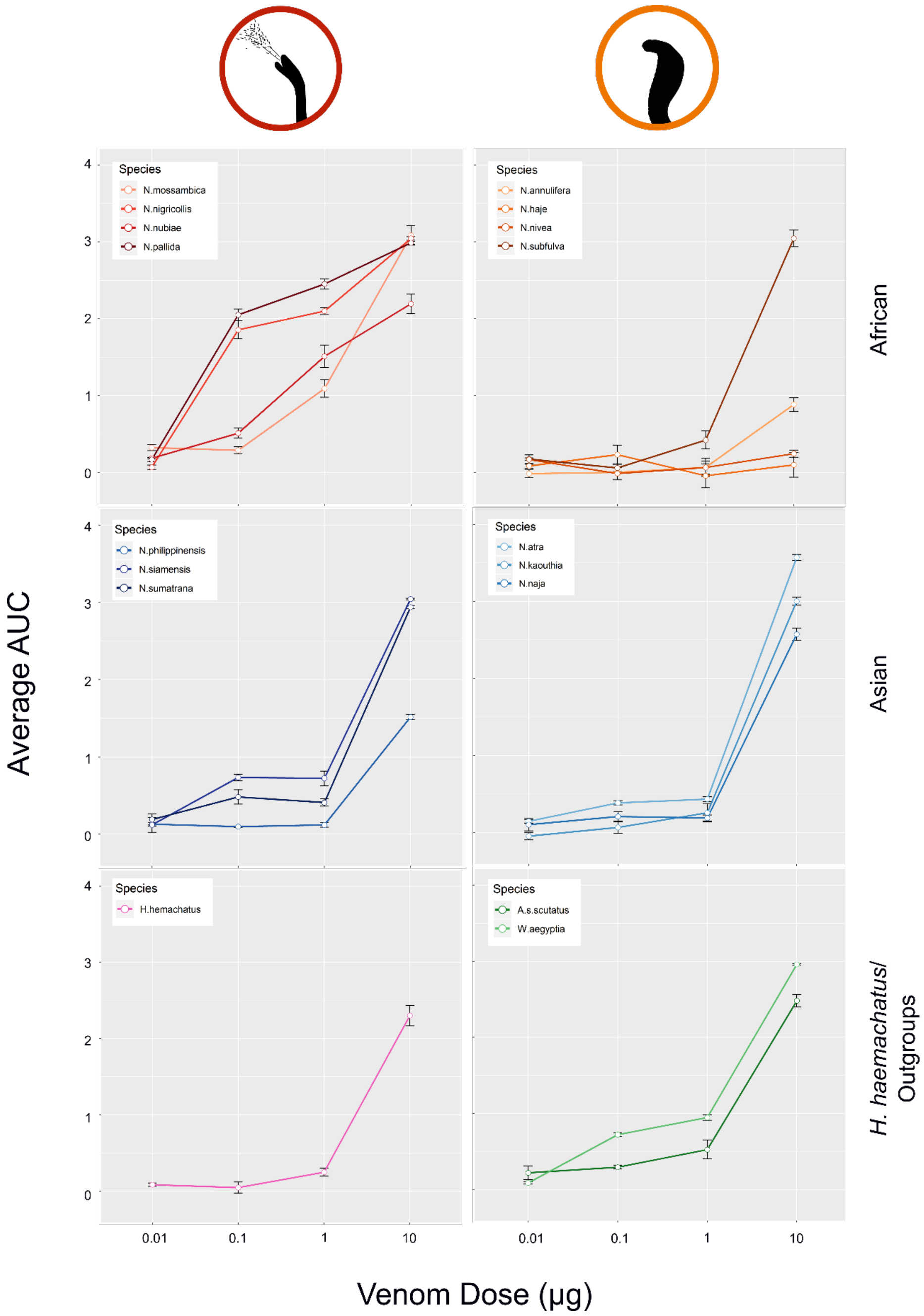


**Fig. S12. Concentration curves displaying the enzymatic $PLA_2$ activity of the various cobra venoms.** The $PLA_2$ activity of each venom was characterized using kinetic measurement of an *in vitro* colorimetric assay (*65*). Venom doses of 0.01 µg, 0.1 µg, 1 µg and 10 µg were quantified in quadruplicate alongside Tris Buffer negative controls. The mean area under the curve (AUC) of the resulting control profile at 10 min was calculated from the four replicates and used to subtract from each individual venom measurement. The mean of the resulting venom data was then plotted, with error bars representing the standard error of the mean of the AUC values at each dose tested.

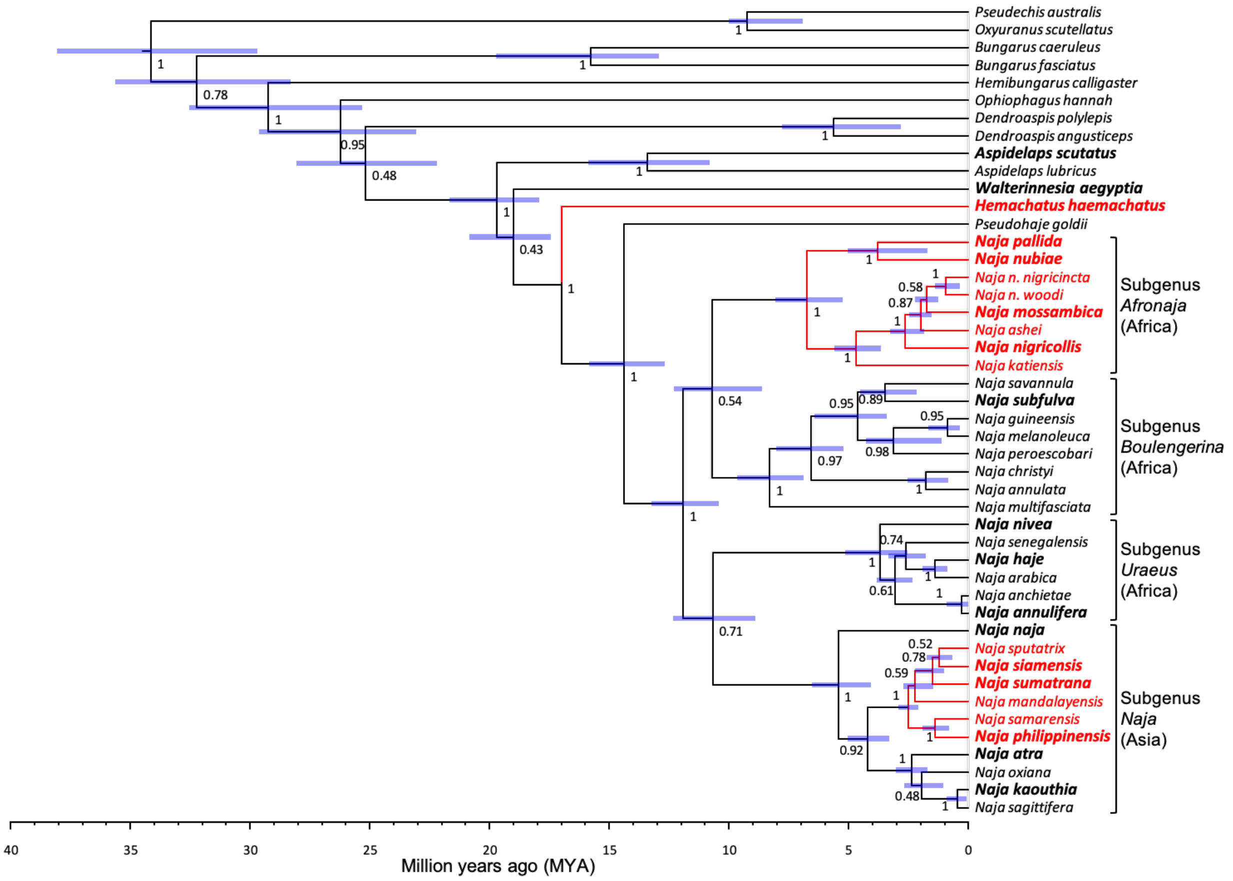


**Fig. S13. Time-calibrated coalescent species tree of cobras and related elapids.** Node labels denote posterior probabilities and blue bars at each node represent age range estimations. Tip labels in bold and enlarged font indicate species included in venom analyses, red tip labels and branches indicate spitting species.

### Supplementary Tables

**Table S1.** The relative representation of phospholipases $A_2$ ($PLA_2$) and three-finger toxins (3FTX) in elapid venom proteomes described in the literature. The data displayed represents the percentage of each toxin family in the whole venom, and note that the method of quantification applied varies among studies.

| **Species** | **3FTX** | | **$PLA_2$** | **Citation** |
|---|---|---|---|---|
| | **CTX** | **Other** | | |
| ***Aipysurus laevis*** | 0 | 25.3 | 71.2 | (*68*) |
| ***Aspidelaps lubricus cowlesi*** | 4.9 | 71.2 | 4.9 | (*16*) |
| ***Aspidelaps lubricus lubricus*** | 2.1 | 75.7 | 5.7 | (*16*) |
| ***Aspidelaps scutatus intermedius*** | 33.7 | 49.0 | 6.1 | (*16*) |
| ***Bungarus caeruleus*** | 0 | 19.0 | 64.5 | (*69*) |
| ***Bungarus candidus*** | 0 | 30.1 | 25.2 | (*70*) |
| ***Bungarus fasciatus*** | 0 | 1.3 | 66.8 | (*71*) |
| ***Bungarus fasciatus*** | 0 | 17.4 | 44.2 | (*70*) |
| ***Bungarus multicinctus*** | 0 | 32.6 | 66.4 | (*72*) |
| ***Bungarus multicinctus*** | 0 | 27.5 | 15.3 | (*71*) |
| ***Bungarus sindanus*** | 0 | 50.3 | 32.6 | (*73*) |
| ***Calliophis intestinalis*** | 15.9* | 5.1 | 43.8 | (*74*) |
| ***Dendroaspis angusticeps*** | 0 | 69.2 | 0 | (*75*) |
| ***Dendroaspis angusticeps*** | 0 | 64.2 | 0 | (*11*) |
| ***Dendroaspis jamesoni jamesoni*** | 0 | 65.5 | 2.2 | (*11*) |
| ***Dendroaspis jamesoni kaimosae*** | 0 | 74.8 | 1.8 | (*11*) |
| ***Dendroaspis polylepis*** | 0 | 31.0 | 0 | (*76*) |
| ***Dendroaspis polylepis*** | 0 | 40.0 | 0 | (*11*) |
| ***Dendroaspis viridis*** | 0 | 76.7 | 0.2 | (*11*) |
| ***Hydrophis curtus*** | 0 | 26.3 | 62.0 | (*77*) |
| ***Hydrophis cyanocinctus*** | 0 | 81.1 | 18.9 | (*78*) |
| ***Hydrophis platurus*** | 0 | 49.9 | 32.9 | (*79*) |
| ***Hydrophis schistosus*** | 0 | 70.5 | 27.5 | (*80*) |
| ***Laticauda colubrina*** | 0.3 | 66.1 | 33.3 | (*81*) |
| ***Micropechis ikaheka*** | 0 | 9.2 | 80.0 | (*82*) |
| ***Micrurus alleni*** | 0 | 77.3 | 10.9 | (*83*) |
| ***Micrurus altirostris*** | 0 | 79.5 | 13.7 | (*84*) |
| ***Micrurus corallinus*** | 34.8 | | 38.5 | (*85*) |
| ***Micrurus dumerilii*** | 0 | 28.1 | 52.0 | (*86*) |
| ***Micrurus frontalis*** | 0 | 42.4 | 49.2 | (*87*) |
| ***Micrurus lemniscatus carvalhoi*** | 2.3 | | 48.6 | (*85*) |
| ***Micrurus lemniscatus lemniscatus*** | 34.3 | | 19.4 | (*85*) |
| ***Micrurus mipartitus*** | 0 | 61.1 | 29.0 | (*88*) |
| ***Micrurus mipartitus*** | 0 | 83.0 | 8.2 | (*88*) |

| | | | | |
|---|---|---|---|---|
| ***Micrurus mosquitensis*** | 0 | 22.5 | 55.6 | (*83*) |
| ***Micrurus nigrocinctus*** | 0 | 38.0 | 48.0 | (*89*) |
| ***Micrurus paraensis*** | 12.3 | | 65.9 | (*85*) |
| ***Micrurus pyrrhocryptus*** | 0 | 27.0 | 17.0 | (*90*) |
| ***Micrurus spixii spixii*** | 9.5 | | 64.0 | (*85*) |
| ***Micrurus spixii spixii*** | 0 | 56.5 | 37.4 | (*87*) |
| ***Micrurus surinamensis*** | 90.1 | | 6.6 | (*85*) |
| ***Micrurus surinamensis*** | 36.8 | | 33.0 | (*85*) |
| ***Micrurus surinamensis*** | 0 | 95.4 | 4.2 | (*87*) |
| ***Micrurus tschudii*** | 0 | 95.2 | 4.1 | (*91*) |
| ***Naja atra*** | 52.9 | 23.5 | 16.8 | (*92*) |
| ***Naja atra*** | 59.4 | 20.5 | 14.0 | (*92*) |
| ***Naja atra*** | 65.3 | 19.0 | 12.2 | (*72*) |
| ***Naja haje*** | 54.0 | 6.0 | 4.0 | (*93*) |
| ***Naja kaouthia*** | 45.7 | 18.0 | 23.5 | (*94*) |
| ***Naja kaouthia*** | 27.6 | 50.7 | 12.2 | (*94*) |
| ***Naja kaouthia*** | 44.9 | 31.5 | 17.4 | (*94*) |
| ***Naja kaouthia*** | 27.9 | 28.6 | 26.9 | (*95*) |
| ***Naja katiensis*** | 62.7 | 4.4 | 29.0 | (*44*) |
| ***Naja melanoleuca***** | 25.2 | 31.9 | 12.9 | (*96*) |
| ***Naja mossambica*** | 67.7 | 1.6 | 27.1 | (*44*) |
| ***Naja naja*** | 63.8 | | 11.4 | (*97*) |
| ***Naja naja*** | 69.3 | 4.8 | 21.4 | (*98*) |
| ***Naja naja*** | 71.6 | 8.9 | 14.0 | (*98*) |
| ***Naja nigricollis*** | 72.8 | 0.5 | 21.9 | (*44*) |
| ***Naja nubiae*** | 58.3 | 12.6 | 26.4 | (*44*) |
| ***Naja pallida*** | 64.9 | 2.8 | 30.1 | (*44*) |
| ***Naja philippinensis*** | 21.3 | 45.3 | 22.9 | (*99*) |
| ***Naja sputatrix*** | 48.1 | 16.1 | 31.2 | (*100*) |
| ***Notechis scutatus*** | 0 | 5.6 | 74.5 | (*101*) |
| ***Ophiophagus hannah*** | 9.0 | 55.2 | 2.8 | (*67*) |
| ***Ophiophagus hannah*** | 0.5 | 42.5 | 4.0 | (*102*) |
| ***Oxyuranus scutellatus*** | 0 | 4.2 | 68.3 | (*103*) |
| ***Oxyuranus scutellatus*** | 0 | 1.5 | 79.4 | (*103*) |
| ***Pseudechis papuanus*** | 0 | 3.1 | 90.2 | (*104*) |
| ***Toxicocalamus longissimus*** | 0 | 92.1 | 6.5 | (*78*) |

* BlastP analysis of the proteomic fragments reported in this study reveals that six of the seven proteins annotated as cytotoxins do not match with cytotoxins as their closest match in the NCBI database. The remaining sequence showed only 44% identity with a cytotoxin.

** This study was published before the recent reclassification of *N. melanoleuca* into five distinct species (*105*). Based on the locality provided in this publication, the venom used in this study was likely sourced from either *N. melanoleuca* or *N. subfulva*.

**Table S2.** A summary of the Phylogenetic Generalized Least Squares (PGLS) tests performed under the scenario of *Naja atra* and *Naja kaouthia* being coded as non-spitting species. All tests were performed using the monophyletic grouping of *Naja* and *Hemachatus*, with the outgroup species of *Walterinnesia aegyptia* and *Aspidelaps scutatus* used to provide additional phylogenetic context. Significant values ($P$ <0.05) are represented by bold red font. Degrees of freedom = 15.

| Test Variable | t | P-value |
|---|---|---|
| **$PLA_2$ proteomic abundance** | **4.24** | **0.0007** |
| **CTX proteomic abundance** | -0.83 | 0.42 |
| **'Other 3FTX' proteomic abundance** | -1.21 | 0.25 |
| **Venom lethality by murine lethal dose 50 ($LD_{50}$)** | 0.86 | 0.40 |
| **Potency of neuronal cell activation ($EC_{50}$)** | **-4.48** | **0.0004** |
| **$PLA_2$ enzymatic activity** | **2.24** | **0.04** |
| **Venom cytotoxicity via HET-CAM assay** | 1.08 | 0.30 |
| **Neuronal cell activation ($EC_{50}$) + $PLA_2$ activity** | **-4.45** | **0.0004** |

**Table S3.** Time-dependent irritation scores described for Luepeke's *in vivo* hen's egg test-chorioallantoic membrane (HET-CAM) assay (*62*).

| Effect | Time (min) | | |
|---|---|---|---|
| | 0.5 | 2.0 | 5.0 |
| **Hyperaemia** | 5 | 3 | 1 |
| **Haemorrhage** | 7 | 5 | 3 |
| **Coagulation** | 9 | 7 | 5 |

**Table S4.** The previously described relationship between the cumulative irritation score of Luepeke's *in vivo* then's egg test-chorioallantoic membrane (HET-CAM) assay (see Table S3) and 'irritation potential' (*62*).

| **Cumulative score** | **Irritation assessment (based on Luepke (*62*))** | **Irritation potential** |
|---|---|---|
| **0.0-0.9** | Practically none | No irritation |
| **1.0-4.9** | Slight | Slight irritation |
| **5.0-8.9** | Moderate | Moderate irritation |
| **9.0-21.0** | Strong | Severe irritation |

**Table S5.** The murine lethality of the elapid snake venoms used in this study represented by intravenous median lethal dose ($LD_{50}$) values and corresponding 95% confidence intervals (CI).

| Species | Group | Lineage | $LD_{50}$ (µg/mouse) | 95% CI |
|---|---|---|---|---|
| ***H. haemachatus*** | Spitter | *H. haemachatus* | 57.96 | 47.28-80.00 |
| ***N. mossambica*** | Spitter | African spitter | 24.48 | 19.68-29.51 |
| ***N. nubiae*** | Spitter | African spitter | 23.65 | 19.17-27.37 |
| ***N. pallida*** | Spitter | African spitter | 9.29 | 3.76-13.23 |
| ***N. nigricollis*** | Spitter | African spitter | 27.49 | 22.55-38.21 |
| ***N. annulifera*** | Non-spitter | African non-spitter | 33.75 | 29.72-37.18 |
| ***N. subfulva*** | Non-spitter | African non-spitter | 5.84 | 3.27-7.47 |
| ***N. nivea*** | Non-spitter | African non-spitter | 15.05 | 10.20-18.70 |
| ***N. haje*** | Non-spitter | African non-spitter | 8.15 | 6.55-9.89 |
| ***N. philippinensis*** | Spitter | Asian spitter | 2.36 | 0.64-4.20 |
| ***N. siamensis*** | Spitter | Asian spitter | 18.43 | 14.02-22.17 |
| ***N. sumatrana*** | Spitter | Asian spitter | 17.43 | 13.29-20.97 |
| ***N. atra*** | Non-spitter | Asian non-spitter | 17.86 | 14.16-27.89 |
| ***N. kaouthia*** | Non-spitter | Asian non-spitter | 6.69 | 3.89-8.54 |
| ***N. naja*** | Non-spitter | Asian non-spitter | 11.1 | 7.90-13.10 |
| ***A. scutatus*** | Non-spitter | Outgroup | 5.75 * | 5.29-6.26 |
| ***W. aegyptia*** | Non-spitter | Outgroup | 15.08 | 11.78-20.68 |

* Previously determined by Whiteley et al. (*16*).

**Table S6.** A summary of the Phylogenetic Generalized Least Squares (PGLS) tests performed under the scenario of *Naja atra* and *Naja kaouthia* being coded as spitting species. All tests were performed using the monophyletic grouping of *Naja* and *Hemachatus*, with the outgroup species of *Walterinnesia aegyptia* and *Aspidelaps scutatus* used to provide additional phylogenetic context. Significant values ($P < 0.05$) are represented by bold red font. Degrees of freedom = 15.

| **Test Variable** | **t** | **P-value** |
|---|---|---|
| **$PLA_2$ proteomic abundance** | **2.86** | **0.01** |
| **CTX proteomic abundance** | -0.57 | 0.58 |
| **'Other 3FTX' proteomic abundance** | -0.76 | 0.46 |
| **Venom lethality by murine lethal dose 50 ($LD_{50}$)** | 1.25 | 0.23 |
| **Potency of neuronal cell activation ($EC_{50}$)** | **-4.59** | **0.0004** |
| **$PLA_2$ enzymatic activity** | 1.80 | 0.09 |
| **Venom cytotoxicity via HET-CAM assay** | **3.29** | **0.004** |
| **Neuronal cell activation ($EC_{50}$) + $PLA_2$ activity** | **-4.57** | **0.0004** |

**Table S7.** The NCBI nucleotide accession numbers associated with the data used to generated the species tree.

| Species | ND4 | cytb | PRLR | NT3 | UBN1 | c-mos | RAG1 |
|---|---|---|---|---|---|---|---|
| *Aspidelaps lubricus* | MT346783-9 | MT346603-9 | MT347202-12 | MT347034-41 | MT347442-53 | MT346956-7 | MT347362-3 |
| *Aspidelaps scutatus* | AY058969, MT346790-8 | AY188007 | MT347214-21 | KX694994, MT347042-8 | MT347454-8 | AY058923, AY187968, KX694796, MT346958-60 | MT347364-6 |
| *Bungarus caeruleus* | AJ830220, MT346799-800 | AJ749305 | MT347222- | MT347049-50 | MT347459 | MT346961-2 | MT347367 |
| *Bungarus fasciatus* | EU547037, MT346801- | EU547086, MT346610 | MT347224- | FJ434090, KX694998, MT347051-2 | MT347460 | AF544732, AY058924, EU366447, MT346963-4 | EU366438, JF357954, MT347368 |
| *Dendroaspis angusticeps* | MT346803-16 | MT346611-24 | MT347226-38 | MT347053-63 | Mt347461-71 | AF544735, MT346965-6 | MT347369 |
| *Dendroaspis polylepis* | MT346817-25 | MT346625-33 | MT347239-42 | MT347064-7 | MT347472-5 | AY058928, FJ387197 | MT347370-1 |
| *Hemachatus haemachatus* | MT346826-38 | MT346634-46 | MT347244-56 | MT347068-78 | MT347476-88 | MT346968-9 | MT347373-4 |
| *Hemibungarus calligaster* | KX130764 | EF137411 | MT347243 | MT347192 | MT347489 | EF137419, MT346967 | MT347372 |
| *Naja (Afronaja) ashei* | GQ359575, MT346839-42 | GQ359493, MT346647-50 | MT347257-8 | MT347079-80 | MT347497-8 | MT346970-1 | MT347375-6 |
| *Naja (Afronaja) katiensis* | GQ359576, MT346843-5 | GQ359494, MT346651-3 | MT347259-60 | MT347081-2 | MT347499-500 | MT346972-3 | MT347377 |
| *Naja (Afronaja) mossambica* | GQ359577, MT346846-53 | AF399747, GQ359495, MT346654-60 | MT347261-3 | MT347083-5 | MT347501-3 | MT346974-5 | MT347378-9 |
| *Naja (Afronaja) nigricincta nigricincta* | MT346854-5 | MT346661-2 | MT347264-5 | MT347086-7 | MT347504-5 | MT346976-7 | MT347380 |
| *Naja (Afronaja) nigricincta woodi* | MT346889-92 | MT346693-6 | MT347290-1 | MT347105-6 | MT347525-6 | MT346984-5 | MT347385 |
| *Naja (Afronaja) nigricollis* | AY713377, MT346856-79 | AF399746, GQ359505, MT346663-85 | MT347266-84 | MT347088-99 | MT347506-19 | MT346978-9 | MT347381 |
| *Naja (Afronaja) nubiae* | GQ359579, MT346880-81 | AF399751, GQ359497, MT346686 | MT347285-7 | MT347100-2 | MT347520-2 | MT346980-1 | MT347382 |
| *Naja (Afronaja) pallida* | GQ359578, MT346882-8 | AF399748, GQ359496, MT346687-92 | MT347288-9 | MT347103-4 | MT347523-4 | MT346982-3 | MT347383-4 |
| *Naja (Boulengerina) annulata* | MT346893-6 | MT346697-700 | MT347292-8 | MT347107-9 | MT347490-2 | AY058925, AY187971, MT346986 | MT347386-7 |
| *Naja (Boulengerina) christyi* | MT346897 | MT346701 | MT347299 | MT347110 | MT347493 | | |
| *Naja (Boulengerina) guineensis* | MH337376-88 | MH337570-82 | MH337474-5, MH337484-95 | MT347111-5 | MH337511-24 | MT346987-8 | MT347388 |
| *Naja (Boulengerina) melanoleuca* | MH337389-402 | MH337583-96 | MH337472-83, MH337496-8, MH337500, MT347301- | MT347116-21 | MH347505-10, MH337525-30, MH337567-8, MT347494-6 | AY611904, MT346989-90 | JF357955, MT347389-90 |

| | | | | | | | |
|---|---|---|---|---|---|---|---|
| *Naja (Boulengerina) multifasciata* | AY058985, MT346898 | AF217837, MT346702 | MT347303 | MT347122 | MT347527 | AY058941 | MT347391 |
| *Naja (Boulengerina) peroescobari* | MH337440 | MH337634 | MH337499 | MT347123-4 | MH337563 | MT346991-2 | MT347392-3 |
| *Naja (Boulengerina) savannula* | MH337403-8 | MH337597-602 | MH337501-4 | MT347125-8 | MH337532-5 | MT346993-4 | MT397394-5 |
| *Naja (Boulengerina) subfulva* | MH337409-21, MH337426-39 | MH337603-33 | MH337441-71 | MT347129-45 | MH337531, MH337537-66 | MT346995-6 | MT347396-7 |
| *Naja (Naja) atra* | MT346899-900 | MT346703-4 | MT347304-5 | MT347146-7 | MT347528-9 | KX694797, MT346997-8 | MT347398-9 |
| *Naja (Naja) kaouthia* | EU624209, MT346901-4 | GQ359507, MT346705-8 | MT347306-8 | MT347148-50 | MT347530-2 | AY058938, MT346999 | MT347400 |
| *Naja (Naja) mandalayensis* | MT346905-6 | MT346709-10 | MT347309-10 | MT347151-2 | MT347533-4 | MT347000 | MT347401-2 |
| *Naja (Naja) naja* | AY713378, MH337375, MT346907-8 | GQ359506, MH337569, MT346711-2 | MT347311-2 | MT347155-6 | MT347535-6 | EU366445, MT347001-2 | EU366432, MT347403 |
| *Naja (Naja) oxiana* | MT346909-10 | MT346713-4 | MT347313-4 | MT347157 | MT347537 | MT347003 | MT347404 |
| *Naja (Naja) philippinensis* | MT346911-5 | MT346911-5 | MT347315-6 | MT347158-9 | MT347538-9 | MT347004-5 | MT347405 |
| *Naja (Naja) sagittifera* | MT346916-7 | MT346916-7 | MT347317 | MT347160 | MT347540 | MT347006-7 | MT347406 |
| *Naja (Naja) samarensis* | MT346918-20 | MT346918-20 | MT347318-9 | MT347161-2 | MT347541-2 | MT347008 | MT347407-8 |
| *Naja (Naja) siamensis* | MT346923-6 | MT346923-6 | MT347320-1 | MT347163-5 | MT347543-5 | MT347009-11 | MT347409-10 |
| *Naja (Naja) sputatrix* | MT346921-2 | MT346921-2 | MT347322 | MT347166 | MT347546 | | |
| *Naja (Naja) sumatrana* | MT346927-34 | MT346927-34 | MT347323-6 | MT347153-4, MT347167-8 | MT347547-51 | MT347012-6 | MT347411-5 |
| *Naja (Uraeus) anchietae* | GQ387085-8 | GQ387085-8 | MT347327-8 | MT347169-70 | MT347552-3 | MT347017 | MT347416 |
| *Naja (Uraeus) annulifera* | GQ359586, GQ387089 | GQ359586, GQ387089 | MT347329-31 | MT347171-2 | MT347554-6 | MT347018 | MT347417-8 |
| *Naja (Uraeus) arabica* | GQ359382, GQ387074-9 | GQ359500, MT346744-8 | MT347332-3 | MT347173-4 | MT347557-8 | MT347019-20 | MT347419-20 |
| *Naja (Uraeus) haje* | GQ359580-1, GQ359583, GQ387062, GQ387064, GQ387066, GQ387068-73 | GQ359498, GQ359501, MT346749-58 | MT347334-9 | MT347175-8 | MT347559-64 | AY611903, MT347021-2 | MT347421-2 |
| *Naja (Uraeus) nivea* | AY058983, MT346935-6 | AF217827, MT346759-60 | MT347340-1 | MT347179-80 | MT347564-6 | AY058939, MT347023 | MT347423-4 |
| *Naja (Uraeus) senegalensis* | GQ359584-5, GQ387080, GQ387083-4 | GQ359502-3, MT346761-3 | MT347342-3 | MT347181-2 | MT347567-8 | MT347024-5 | MT347425 |
| *Ophiophagus hannah* | AY058984, EU921899, MT346937-46 | AF217842, EU921899, MT346764-73 | MT347344-53 | MT347183-91 | MT347569-74 | AY058940, KX694798, MT347026 | MT347426-34 |
| *Oxyuranus scutellatus* | EU547006, MT346947 | EU547051, MT346774 | MT347354-5 | MT347193-4 | MT347575-6 | EU546916, MT347027-8 | EU546877, MT347435-6 |
| *Pseudechis australis* | AJ83071, MT346948-50 | EU547046, MT346775-7 | KX981746-7, MT347356-7 | MT347195-7 | MT347577-80 | EU546912, MT347029 | EU546873, MT347437 |
| *Pseudohaje goldii* | MT346951-2 | MT346778-9 | MT347358 | MT347198 | MT347581 | MT347030 | MT347438 |
| *Walterinnesia aegyptia* | AY058988, MT346953-5 | AF217838, MT346780-2 | MT347359-61 | MT347199-201 | MT347582-4 | AY058943, MT347031-3 | MT347439-41 |

**Table S8.** The snake species, localities of origin, and number of specimens that contributed to the venom pools used in venom proteomic analyses and murine lethality tests.

| Species | Locality | Number of individual snakes contributing to the venom pool |
|---|---|---|
| ***Hemachatus haemachatus*** | Captive bred | 2 |
| ***Naja annulifera*** | Captive bred | 2 |
| ***Naja atra*** | Captive bred | 2 |
| ***Naja haje*** | Uganda | 6 |
| ***Naja kaouthia*** | Captive bred | 3 |
| ***Naja mossambica*** | Tanzania | 4 |
| ***Naja naja*** | Captive bred | 2 |
| ***Naja nigricollis*** | Nigeria | 3 |
| ***Naja nivea*** | South Africa | 3 |
| ***Naja nubiae*** | Captive bred | 3 |
| ***Naja pallida*** | Tanzania | 2 |
| ***Naja philippinensis*** | Captive bred | 2 |
| ***Naja siamensis*** | Captive bred | 2 |
| ***Naja subfulva*** | Cameroon | 2 |
| ***Naja sumatrana*** | Captive bred | 1 |
| ***Walterinnesia aegyptia*** | Captive bred | 2 |

**Table S9.** PCR primers and typical thermocycling conditions for the genes sequenced for the phylogenetic analysis. The denaturing step involved 30-45 s at 94°C and the extension step 1 min at 72°C, with a final extension step of 5 min at 72°C followed by cooling to 4°C for 15 min.

| **Gene** | **Sense** | **Name** | **Sequence** | **Citation** | **Annealing temp.** | **Cycles** |
|---|---|---|---|---|---|---|
| **CytB** | Forward | Gludg | 5'– TGACTTGAARAACCAYCGTTG – 3' | (*106*) | 47°C | 39 |
| | Reverse | ATRCB3 | 5'– TGAGAAGTTTTCYGGGTCRTT – 3' | (*107*) | | |
| | | H16064 | 5'– CTTTGGTTTACAAGAACAATGCTTTA – 3' | (*108*) | | |
| **ND4** | Forward | NADH4 | 5'– CACCTATGACTACCAAAAGCTCATGTAGAAGC – 3' | (*109*) | 57°C | 39 |
| | Reverse | H12763V | 5'– TTCTATCACTTGGATTTGCACCA – 3' | (*109*) | | |
| **C-MOS** | Forward | AV CMOSF | 5'– AAGCACATCAAGGATTCGTCG – 3' | (*110*) | 60.5°C | 44 |
| | Reverse | AV CMOSR | 5'– TCTGCCTTGGGTGTGATTTTCT – 3' | | | |
| | Forward | G303 | 5'– ATTATGCCATCMCCTMTTCC – 3' | (*111*) | 57°C | 35 |
| | Reverse | G708 | 5'– GCTACATCAGCTCTCCARCA – 3' | | | |
| **NT3** | Forward | NTF3 F1 | 5'– ATGTCCAATCTTGTTTTATGTGATATTT – 3' | (*112*) | 42°C | 39 |
| | Reverse | NTF3 R1 | 5'– ACRAGTTTRTTGTTYTCTGAAGTC – 3' | | | |
| **PRLR** | Forward | PRLR F1 | 5'– GACARYGARGACCAGCAACTRATGCC – 3' | (*112*) | 48°C | 39 |
| | Reverse | PRLR F3 | 5'– GACYTTGTGRACTTCYACRTAATCCAT – 3' | | | |
| **RAG1** | Forward | AV RAG1F | 5'– AAATGTGACAGGGTCTCT – 3' | (*110*) | 59°C | 44 |
| | Reverse | AV RAG1R | 5'– GGGCATCTCAAAACCAAATTGT – 3' | | | |
| **UBN1** | Forward | BaUBN F | 5'– CCTCTGGTTACTCAGCAGCA – 3' | (*113*) | 40°C | 39 |
| | Reverse | BaUBN R | 5'– ATTGGCCACTCCTTGTGTTC – 3' | | | |

**Table S10.** The snake species, localities of origin, and number of specimens that contributed to the venom pools used in the hen's egg test-chorioallantoic membrane (HET-CAM) assay.

| Species | Locality | Number of individual snakes contributing to the venom pool | Same venom sample as that used for proteomic analysis? |
|---|---|---|---|
| ***Naja kaouthia*** | Captive bred | 1 | No |
| ***Naja mossambica*** | Captive bred | 1 | No |
| ***Naja nivea*** | Captive bred | 1 | No |
| ***Naja atra*** | Captive bred | 1 | No |
| ***Naja sumatrana*** | Captive bred | 1 | No |
| ***Naja siamensis*** | Captive bred | 1 | No |
| ***Naja pallida*** | Captive bred | 1 | No |
| ***Naja annulifera*** | Captive bred | 1 | No |
| ***Naja haje*** | Captive bred | 1 | No |
| ***Naja subfulva*** | Captive bred | 1 | No |
| ***Naja naja*** | Captive bred | 1 | No |
| ***Naja nubiae*** | Captive bred | 1 | No |
| ***Walterinnesia aegyptia*** | Captive bred | 2 | Yes |
| ***Hemachatus haemachatus*** | Captive bred | 2 | Yes |
| ***Naja philippinensis*** | Captive bred | 2 | Yes |
| ***Naja nigricollis*** | Nigeria | 3 | Yes |
| ***Aspidelaps lubricus cowlesi*** | Captive bred | 1 | N/A (proteomics previously performed) |

### Supplementary Data Files

#### Data S1.

Top-down MS analyses of the venom proteomes described in this study: including African and Asian *Naja* species, the desert black snake (*Walterinnesia aegyptia*), and the rinkhals (*Hemachatus haemachatus*), as summarized in Fig. 1A and Fig. S2.

#### Data S2.

Nucleotide sequence alignment for the three-finger toxin (3FTX) family, extracted from venom gland transcriptomes from *Naja* spp., the desert black snake (*Walterinnesia aegyptia*) and the rinkhals (*Hemachatus haemachatus*) described in this study, along with data from related elapid snake species (*Aspidelaps scutatus intermedius* (*16*), *Bungarus flaviceps* (*52*), *Bungarus multicinctus* (52), *Dendroaspis* spp. (*11*), *Micrurus fulvius* (*55*) and *Ophiophagus hannah* (53, 54)) sourced from previous, aligned against a sequence from the outgroup species (*Python regius*; NCBI accession number GBIC00000000, from (*57*)).

#### Data S3.

Nucleotide sequence alignment for the cytotoxin (CTX) subset of the three-finger toxin (3FTX) family, extracted from venom gland transcriptomes from *Naja* spp., the rinkhals (*Hemachatus haemachatus*), the shield-nosed cobra (*Aspidelaps scutatus intermedius*, from (*16*)) and the king cobra (*Ophiophagus hannah*, from (*54*)), aligned against a sequence from the outgroup elapid, the red-headed krait (*Bungarus flaviceps*, NCBI accession number GU190795, from (*52*)).

### Supplementary References


33. D. A. Dmitriev, R. A. Rakitov, Decoding of Superimposed Traces Produced by Direct Sequencing of Heterozygous Indels. *PLoS Comput. Biol.* **4**, e1000113 (2008).

34. M. Stephens, P. Scheet, Accounting for Decay of Linkage Disequilibrium in Haplotype Inference and Missing-data Imputation. *Am. J. Hum. Genet.* **76**, 449–462 (2005).

35. M. Stephens, N. J. Smith, P. Donnelly, A New Statistical Method for Haplotype Reconstruction from Population Data. *Am. J. Hum. Genet.* **68**, 978–989 (2001).

36. J. F. Flot, Seqphase: A Web Tool for Interconverting Phase Input/Output Files and Fasta Sequence Alignments. *Mol. Ecol. Resour.* **10**, 162–166 (2010).

37. A. J. Drummond, M. A. Suchard, D. Xie, A. Rambaut, Bayesian Phylogenetics with BEAUti and the BEAST 1.7. *Mol. Biol. Evol.* **29**, 1969–1973 (2012).

38. J. J. Head, K. Mahlow, J. Müller, Fossil Calibration Dates for Molecular Phylogenetic Analysis of Snakes 2: Caenophidia, Colubroidea, Elapoidea, Colubridae. *Palaeontol. Electron.* **19**, 1–21 (2016).

39. D. Pla, L. Sanz, G. Whiteley, S. C. Wagstaff, R. A. Harrison, N. R. Casewell, J. J. Calvete, What Killed Karl Patterson Schmidt? Combined Venom Gland Transcriptomic, Venomic and Antivenomic Analysis of the South African Green Tree Snake (the Boomslang), *Dispholidus typus*. *Biochim. Biophys. Acta - Gen. Subj.* **1861**, 814–823 (2017).

40. J. Archer, G. Whiteley, N. R. Casewell, R. A. Harrison, S. C. Wagstaff, VTBuilder: a Tool for the Assembly of Multi Isoform Transcriptomes. *BMC Bioinformatics*. **15**, 389 (2014).

41. A. Conesa, S. Götz, J. M. García-Gómez, J. Terol, M. Talón, M. Robles, Blast2GO: A Universal Tool for Annotation, Visualization and Analysis in Functional Genomics Research. *Bioinformatics*. **21**, 3674–3676 (2005).

42. S. Kumar, G. Stecher, K. Tamura, MEGA7: Molecular Evolutionary Genetics Analysis Version 7.0 for Bigger Datasets. *Mol. Biol. Evol.* **33**, 1870-1874 (2016).

43. R. C. Edgar, MUSCLE: Multiple Sequence Alignment with High Accuracy and High Throughput. *Nucleic Acids Res.* **32**, 1792–1797 (2004).

44. D. Petras, L. Sanz, Á. Segura, M. Herrera, M. Villalta, D. Solano, M. Vargas, G. León, D. A. Warrell, R. D. G. Theakston, R. A. Harrison, N. Durfa, A. Nasidi, J. M. Gutiérrez, J. J. Calvete, Snake venomics of African Spitting Cobras: Toxin Composition and Assessment of Congeneric Cross-reactivity of the Pan-African EchiTAb-Plus-ICP Antivenom by Antivenomics and Neutralization Approaches. *J. Proteome Res.* **10**, 1266–1280 (2011).

45. D. Pla, D. Petras, A. J. Saviola, C. M. Modahl, L. Sanz, A. Pérez, E. Juárez, S. Frietze, P. C. Dorrestein, S. P. Mackessy, J. J. Calvete, Transcriptomics-guided Bottom-up and Top-down Venomics of Neonate and Adult Specimens of the Arboreal Rear-fanged Brown Treesnake, *Boiga irregularis*, from Guam. *J. Proteomics*. **174**, 71–84 (2018).

46. Q. Kou, L. Xun, X. Liu, TopPIC: A Software Tool for Top-down Mass Spectrometry-Based

Proteoform Identification and Characterization. *Bioinformatics*. **32**, 3495–3497 (2016).

47. L. J. Revell, phytools: An R Package for Phylogenetic Comparative Biology (and other things). *Methods Ecol. Evol.* **3**, 217–223 (2012).

48. D. Orme, R. Freckleton, G. Thomas, T. Petzoldt, S. Fritz, N. I. and W. Pearse, caper: Comparative Analyses of Phylogenetics and Evolution in R (2018), (available at https://cran.r-project.org/package=caper).

49. RStudio, RStudio: Integrated Development for R. RSudio, Inc., Boston, MA (2016), (available at http://www.rstudio.com/).

50. D. T. Jones, W. R. Taylor, J. M. Thornton, The Rapid Generation of Mutation Data matrices from protein sequences. *Comput. Appl. Biosci.* **8**, 275–282 (1992).

51. M. Maechler, P. Rousseeuw, A. Struyf, M. Hubert, K. Hornik, cluster: Cluster Analysis Basics and Extensions. R package version 2.1.0 (2019).

52. A. S. Siang, R. Doley, F. J. Vonk, R. M. Kini, Transcriptomic Analysis of the Venom Gland of the Red-headed Krait (*Bungarus flaviceps*) Using Expressed Sequence Tags. *BMC Mol. Biol.* **11**, 24 (2010).

53. Y. Jiang, Y. Li, W. Lee, X. Xu, Y. Zhang, R. Zhao, Y. Zhang, W. Wang, Venom Gland Transcriptomes of Two Elapid Snakes (*Bungarus multicinctus* and *Naja atra*) and Evolution of Toxin Genes. *BMC Genomics*. **12**, 1 (2011).

54. Li,J., Zhang, J. H., Liu, K. Xu, Novel Genes Encoding Six Kinds of Three-finger Toxins in *Ophiophagus hannah* (King Cobra) and Function Characterization of Two Recombinant Long-chain Neurotoxins. *Biochem. J.* **398**, 233–242 (2006).

55. M. J. Margres, K. Aronow, J. Loyacano, D. R. Rokyta, The Venom-gland Transcriptome of the Eastern Coral Snake (*Micrurus fulvius*) Reveals High Venom Complexity in the Intragenomic Evolution of Venoms. *BMC Genomics*. **14**, 531 (2013).

56. F. J. Vonk, N. R. Casewell, C. V Henkel, A. M. Heimberg, H. J. Jansen, R. J. R. McCleary, H. M. E. Kerkkamp, R. A. Vos, I. Guerreiro, J. J. Calvete, W. Wüster, A. E. Woods, J. M. Logan, R. A. Harrison, T. A. Castoe, A. P. J. de Koning, D. D. Pollock, M. Yandell, D. Calderon, C. Renjifo, R. B. Currier, D. Salgado, D. Pla, L. Sanz, A. S. Hyder, J. M. C. Ribeiro, J. W. Arntzen, G. E. E. J. M. van den Thillart, M. Boetzer, W. Pirovano, R. P. Dirks, H. P. Spaink, D. Duboule, E. McGlinn, R. M. Kini, M. K. Richardson, The King Cobra Genome Reveals Dynamic Gene Evolution and Adaptation in the Snake Venom System. *Proc. Natl. Acad. Sci. U. S. A.* **110**, 20651–20656 (2013).

57. A. D. Hargreaves, M. T. Swain, D. W. Logan, J. F. Mulley, Testing the Toxicofera: Comparative Transcriptomics Casts Doubt on the Single, Early Evolution of the Reptile Venom System. *Toxicon*. **92**, 140–156 (2014).

58. D. Darriba, G. L. Taboada, R. Doallo, D. Posada, jModelTest 2: more Models, New Heuristics and Parallel Computing. *Nat. Methods*. **9**, 772–772 (2012).

59. D. Posada, jModelTest: Phylogenetic Model Averaging. *Mol. Biol. Evol.* **25**, 1253–1256

(2008).

60. F. Ronquist, M. Teslenko, P. Van Der Mark, D. L. Ayres, A. Darling, S. Höhna, B. Larget, L. Liu, M. A. Suchard, J. P. Huelsenbeck, Mrbayes 3.2: Efficient Bayesian Phylogenetic Inference and Model Choice Across a Large Model Space. *Syst. Biol.* **61**, 539–542 (2012).

61. M. A. Miller, W. Pfeiffer, T. Schwartz, Creating the CIPRES Science Gateway for Inference of Large Phylogenetic Trees. *2010 Gatew. Comput. Environ. Work. (GCE),* New Orleans, LA, pp. 1-8 (2010).

62. N. P. Luepke, Hen's Egg Chorioallantoic Membrane Test for Irritation Potential. *Food Chem. Toxicol.* **23**, 287–291 (1985).

63. I. Vetter, R. J. Lewis, Characterization of Endogenous Calcium Responses in Neuronal Cell Lines. *Biochem. Pharmacol.* **79**, 908–920 (2010).

64. J. E. Hale, J. P. Butler, V. Gelfanova, J. S. You, M. D. Knierman, A Simplified Procedure for the Reduction and Alkylation of Cysteine Residues in Proteins Prior to Proteolytic Digestion and Mass Spectral Analysis. *Anal. Biochem.* **333**, 174–181 (2004).

65. K. B. M. Still, J. Slagboom, S. Kidwai, C. Xie, B. Eisses, J. Freek, Development of High-throughput Screening Assays for Profiling Snake Venom Phospholipase A 2 Activity after High-resolution Chromatographic Fractionation *bioRxiv.* (2020). doi: 10.1101/2020.01.20.912758.

66. R. D. G. Theakston, H. A. Reid, Development of Simple Standard Assay Procedures for the Characterization of Snake Venoms. **61**, 949–956 (1983).

67. D. Petras, P. Heiss, R. D. Süssmuth, J. J. Calvete, Venom Proteomics of Indonesian King Cobra, *Ophiophagus hannah*: Integrating Top-down and Bottom-up approaches. *J. Proteome Res.* **14**, 2539–2556 (2015).

68. A. H. Laustsen, J. M. Gutiérrez, A. R. Rasmussen, M. Engmark, P. Gravlund, K. L. Sanders, B. Lohse, B. Lomonte, Danger in the Reef: Proteome, Toxicity, and Neutralization of the Venom of the Olive Sea Snake, *Aipysurus laevis*. *Toxicon*. **107**, 187–196 (2015).

69. A. M. F. Oh, C. H. Tan, G. C. Ariaranee, N. Quraishi, N. H. Tan, Venomics of *Bungarus caeruleus* (Indian Krait): Comparable Venom Profiles, Variable Immunoreactivities Among Specimens from Sri Lanka, India and Pakistan. *J. Proteomics*. **164**, 1–18 (2017).

70. M. R. A. Rusmili, T. T. Yee, M. R. Mustafa, W. C. Hodgson, I. Othman, Proteomic Characterization and Comparison of Malaysian Bungarus candidus and *Bungarus fasciatus* Venoms. *J. Proteomics*. **110**, 129–144 (2014).

71. R. H. Ziganshin, S. I. Kovalchuk, G. P. Arapidi, V. G. Starkov, A. N. Hoang, T. T. Thi Nguyen, K. C. Nguyen, B. B. Shoibonov, V. I. Tsetlin, Y. N. Utkin, Quantitative Proteomic Analysis of Vietnamese Krait Venoms: Neurotoxins are the Major Components in *Bungarus multicinctus* and Phospholipases A2 in *Bungarus fasciatus*. *Toxicon*. **107**, 197–209 (2015).

72. L. L. Shan, J. F. Gao, Y. X. Zhang, S. S. Shen, Y. He, J. Wang, X. M. Ma, X. Ji, Proteomic

Characterization and Comparison of Venoms from Two Elapid Snakes (*Bungarus multicinctus* and *Naja atra*) from China. *J. Proteomics*. **138**, 83–94 (2016).

73. A. M. F. Oh, C. H. Tan, K. Y. Tan, N. H. Quraishi, N. H. Tan, Venom Proteome of *Bungarus sindanus* (Sind Krait) from Pakistan and in vivo Cross-neutralization of Toxicity using an Indian Polyvalent Antivenom. *J. Proteomics*. **193**, 243–254 (2019).

74. K. Y. Tan, J. L. Liew, N. H. Tan, E. S. H. Quah, A. K. Ismail, C. H. Tan, Unlocking the Secrets of Banded Coral Snake (*Calliophis intestinalis*, Malaysia): A Venom with Proteome Novelty, Low Toxicity and Distinct Antigenicity. *J. Proteomics*. **192**, 246–257 (2019).

75. L. P. Lauridsen, A. H. Laustsen, B. Lomonte, J. M. Gutiérrez, Toxicovenomics and Antivenom Profiling of the Eastern Green Mamba Snake (*Dendroaspis angusticeps*). *J. Proteomics*. **136**, 248–261 (2016).

76. A. H. Laustsen, B. Lomonte, B. Lohse, J. Fernández, J. M. Gutiérrez, Unveiling the Nature of Black Mamba (*Dendroaspis polylepis*) Venom through Venomics and Antivenom Immunoprofiling: Identification of Key Toxin Targets for Antivenom Development. *J. Proteomics*. **119**, 126–142 (2015).

77. C. H. Tan, K. Y. Tan, T. S. Ng, S. M. Sim, N. H. Tan, Venom Proteome of Spine-bellied Sea Snake (*Hydrophis curtus*) from Penang, Malaysia: Toxicity Correlation, Immunoprofiling and Cross-neutralization by Sea Snake Antivenom. *Toxins (Basel)*. **11**, 1–19 (2019).

78. J. J. Calvete, P. Ghezellou, O. Paiva, T. Matainaho, A. Ghassempour, H. Goudarzi, F. Kraus, L. Sanz, D. J. Williams, Snake Venomics of Two Poorly Known Hydrophiinae: Comparative Proteomics of the Venoms of Terrestrial *Toxicocalamus longissimus* and Marine *Hydrophis cyanocinctus*. *J. Proteomics*. **75**, 4091–4101 (2012).

79. B. Lomonte, D. Pla, M. Sasa, W. C. Tsai, A. Solórzano, J. M. Ureña-Díaz, M. L. Fernández-Montes, D. Mora-Obando, L. Sanz, J. M. Gutiérrez, J. J. Calvete, Two Color Morphs of the Pelagic Yellow-bellied Sea Snake, *Pelamis platura*, from Different Locations of Costa Rica: Snake Venomics, Toxicity, and Neutralization by Antivenom. *J. Proteomics*. **103**, 137–152 (2014).

80. C. H. Tan, K. Y. Tan, S. E. Lim, N. H. Tan, Venomics of the beaked sea snake, *Hydrophis schistosus*: A Minimalist Toxin Arsenal and its Cross-neutralization by Heterologous Antivenoms. *J. Proteomics*. **126**, 121–130 (2015).

81. C. H. Tan, K. Y. Wong, K. Y. Tan, N. H. Tan, Venom Proteome of the Yellow-lipped Sea Krait, *Laticauda colubrina* from Bali: Insights into Subvenomic Diversity, Venom Antigenicity and Cross-neutralization by Antivenom. *J. Proteomics*. **166**, 48–58 (2017).

82. O. Paiva, D. Pla, C. E. Wright, M. Beutler, L. Sanz, J. M. Gutiérrez, D. J. Williams, J. J. Calvete, Combined Venom Gland cDNA Sequencing and Venomics of the New Guinea Small-eyed snake, *Micropechis ikaheka*. *J. Proteomics*. **110**, 209–229 (2014).

83. J. Fernández, N. Vargas-Vargas, D. Pla, M. Sasa, P. Rey-Suárez, L. Sanz, J. M. Gutiérrez, J. J. Calvete, B. Lomonte, Snake Venomics of *Micrurus alleni* and *Micrurus mosquitensis*

from the Caribbean Region of Costa Rica Reveals Two Divergent Compositional Patterns in New World Elapids. *Toxicon*. **107**, 217–233 (2015).

84. C. Corrêa-Netto, I. de L. M. Junqueira-de-Azevedo, D. A. Silva, P. L. Ho, M. Leitão-de-Araújo, M. L. M. Alves, L. Sanz, D. Foguel, R. B. Zingali, J. J. Calvete, Snake Venomics and Venom Gland Transcriptomic Analysis of Brazilian Coral Snakes, *Micrurus altirostris* and *M. corallinus*. *J. Proteomics*. **74**, 1795–1809 (2011).

85. S. D. Aird, N. J. da Silva, L. Qiu, A. Villar-Briones, V. A. Saddi, M. P. de C. Telles, M. L. Grau, A. S. Mikheyev, Coralsnake venomics: Analyses of Venom Gland Transcriptomes and Proteomes of Six Brazilian Taxa. *Toxins (Basel)*. **9**, 1–64 (2017).

86. P. Rey-Suárez, V. Núñez, J. Fernández, B. Lomonte, Integrative Characterization of the Venom of the Coral Snake *Micrurus dumerilii* (Elapidae) from Colombia: Proteome, Toxicity, and Cross-neutralization by Antivenom. *J. Proteomics*. **136**, 262–273 (2016).

87. L. Sanz, L. N. de Freitas-Lima, S. Quesada-Bernat, V. K. Graça-de-Souza, A. M. Soares, J. J. Calvete, C. A. S. Caldeira, Comparative Venomics of Brazilian Coral Snakes: *Micrurus frontalis*, *Micrurus spixii spixii*, and *Micrurus surinamensis*. *Toxicon*. **166**, 39–45 (2019).

88. P. Rey-Suárez, V. Núñez, J. M. Gutiérrez, B. Lomonte, Proteomic and Biological Characterization of the Venom of the Redtail Coral snake, *Micrurus mipartitus* (Elapidae), from Colombia and Costa Rica. *J. Proteomics*. **75**, 655–667 (2011).

89. J. Fernández, A. Alape-Girón, Y. Angulo, L. Sanz, J. M. Gutiérrez, J. J. Calvete, B. Lomonte, Venomic and Antivenomic Analyses of the Central American Coral Snake, *Micrurus nigrocinctus* (Elapidae). *J. Proteome Res.* **10**, 1816–1827 (2011).

90. T. Olamendi-Portugal, C. V. F. Batista, M. Pedraza-Escalona, R. Restano-Cassulini, F. Z. Zamudio, M. Benard-Valle, A. Rafael de Roodt, L. D. Possani, New Insights into the Proteomic Characterization of the Coral Snake *Micrurus pyrrhocryptus* venom. *Toxicon*. **153**, 23–31 (2018).

91. L. Sanz, D. Pla, A. Pérez, Y. Rodríguez, A. Zavaleta, M. Salas, B. Lomonte, J. J. Calvete, Venomic Analysis of the Poorly Studied Desert Coral Snake, *Micrurus tschudii tschudii*, Supports the 3FTx/PLA2 Dichotomy across *Micrurus* venoms. *Toxins (Basel)*. **8**, 9–12 (2016).

92. H.-W. Huang, B.-S. Liu, K.-Y. Chien, L.-C. Chiang, S.-Y. Huang, W.-C. Sung, W.-G. Wu, Cobra Venom Proteome and Glycome Determined from Individual Snakes of *Naja atra* Reveal Medically Important Dynamic Range and Systematic Geographic Variation. *J. Proteomics*. **128**, 92–104 (2015).

93. I. Malih, M. R. Ahmad rusmili, T. Y. Tee, R. Saile, N. Ghalim, I. Othman, Proteomic Analysis of Moroccan Cobra *Naja haje legionis* Venom Using Tandem Mass Spectrometry. *J. Proteomics*. **96**, 240–252 (2014).

94. K. Y. Tan, C. H. Tan, S. Y. Fung, N. H. Tan, Venomics, Lethality and Neutralization of *Naja kaouthia* (Monocled Cobra) Venoms from three Different Geographical Regions of Southeast Asia. *J. Proteomics*. **120**, 105–125 (2015).

95. N. Xu, H. Y. Zhao, Y. Yin, S. S. Shen, L. L. Shan, C. X. Chen, Y. X. Zhang, J. F. Gao, X. Ji, Combined Venomics, Antivenomics and Venom Gland Transcriptome Analysis of the Monocoled Cobra (*Naja kaouthia*) from China. *J. Proteomics*. **159**, 19–31 (2017).

96. L. P. Lauridsen, A. H. Laustsen, B. Lomonte, J. M. Gutiérrez, Exploring the Venom of the Forest Cobra Snake: Toxicovenomics and Antivenom Profiling of *Naja melanoleuca*. *J. Proteomics*. **150**, 98–108 (2017).

97. S. Dutta, A. Chanda, B. Kalita, T. Islam, A. Patra, A. K. Mukherjee, Proteomic Analysis to Unravel the Complex Venom Proteome of Eastern India *Naja naja*: Correlation of Venom Composition with its Biochemical and Pharmacological Properties. *J. Proteomics*. **156**, 29–39 (2017).

98. K. Sintiprungrat, K. Watcharatanyatip, W. D. S. T. Senevirathne, P. Chaisuriya, D. Chokchaichamnankit, C. Srisomsap, K. Ratanabanangkoon, A Comparative Study of Venomics of *Naja naja* from India and Sri Lanka, Clinical Manifestations and Antivenomics of an Indian Polyspecific Antivenom. *J. Proteomics*. **132**, 131–143 (2016).

99. C. H. Tan, K. Y. Wong, H. P. Chong, N. H. Tan, K. Y. Tan, Proteomic Insights into Short Neurotoxin-driven, Highly Neurotoxic Venom of Philippine Cobra (*Naja philippinensis)* and Toxicity Correlation of Cobra Envenomation in Asia. *J. Proteomics*. **206**, 103418 (2019).

100. N. H. Tan, K. Y. Wong, C. H. Tan, Venomics of *Naja sputatrix*, the Javan Spitting Cobra: A Short Neurotoxin-driven Venom Needing Improved Antivenom Neutralization. *J. Proteomics*. **157**, 18–32 (2017).

101. C. H. Tan, K. Y. Tan, N. H. Tan, Revisiting *Notechis scutatus* Venom: On Shotgun Proteomics and Neutralization by the "Bivalent" Sea Snake Antivenom. *J. Proteomics*. **144**, 33–38 (2016).

102. C. H. Tan, K. Y. Tan, S. Y. Fung, N. H. Tan, Venom-gland Transcriptome and Vnom Proteome of the Malaysian King Cobra (*Ophiophagus hannah*). *BMC Genomics*. **16**, 687 (2015).

103. M. Herrera, J. Fernández, M. Vargas, M. Villalta, Á. Segura, G. León, Y. Angulo, O. Paiva, T. Matainaho, S. D. Jensen, K. D. Winkel, J. J. Calvete, D. J. Williams, J. M. Gutiérrez, Comparative Proteomic Analysis of the Venom of the Taipan Snake, *Oxyuranus scutellatus*, from Papua New Guinea and Australia: Role of Neurotoxic and Procoagulant Effects in Venom Toxicity. *J. Proteomics*. **75**, 2128–2140 (2012).

104. D. Pla, B. W. Bande, R. E. Welton, O. K. Paiva, L. Sanz, Á. Segura, C. E. Wright, J. J. Calvete, J. M. Gutiérrez, D. J. Williams, Proteomics and Antivenomics of Papuan Black Snake (*Pseudechis papuanus*) Venom with Analysis of its Toxicological Profile and the Preclinical Efficacy of Australian Antivenoms. *J. Proteomics*. **150**, 201–215 (2017).

105. W. Wüster, L. Chirio, J. F. Trape, I. Ineich, K. Jackson, E. Greenbaum, C. Barron, C. Kusamba, Z. T. Nagy, R. Storey, C. Hall, C. E. Wüster, A. Barlow, D. G. Broadley, Integration of Nuclear and Mitochondrial Gene Sequences and Morphology Reveals Unexpected Diversity in the Forest Cobra (*Naja melanoleuca*) Species Complex in Central

and West Africa (Serpentes: Elapidae). *Zootaxa*. **4455**, 68–98 (2018).

106. S. R. Palumbi, in *Molecular Systematics, 2nd edition*, D. M. Hillis, C. Moritz, B. K. Mable, Eds. (Sinauer, Sunderland, Massachusetts, 1996), pp. 205–247.

107. M. B. Harvey, D. G. Barker, L. K. Ammerman, P. T. Chippindale, Systematics of Pythons of the *Morelia amethistina* Complex (Serpentes: Boidae) with the Description of Three New Species. *Herpetol. Monogr.* **14**, 139 (2000).

108. F. T. Burbrink, R. Lawson, J. B. Slowinski, Mitochondrial DNA Phylogeography of the Polytypic North American Rat Snake (*Elaphe obsoleta*): A Critique of the Subspecies Concept. *Evolution (N. Y).* **54**, 2107–2118 (2000).

109. E. Arévalo, J. W. Sites, S. K. Davis, E. Arévalo, Mitochondrial DNA Sequence Divergence and Phylogenetic Relationships Among Eight Chromosome Races of the *sceloporus grammicus* Complex (*phrynosomatidae*) in Central Mexico. *Syst. Biol.* **43**, 387–418 (1994).

110. A. von Plettenberg Laing, A Multilocus Phylogeny of the Cobra Clade Elapids. MSc Thesis, Bangor University, Bangor, UK (2019).

111. A. F. Hugall, M. S. Y. Lee, Molecular Claims of Gondwanan Age for Australian Agamid Lizards are Untenable. *Mol. Biol. Evol.* **21**, 2102–2110 (2004).

112. T. M. Townsend, R. E. Alegre, S. T. Kelley, J. J. Wiens, T. W. Reeder, Rapid Development of Multiple Nuclear Loci for Phylogenetic Analysis using Genomic Resources: An Example from Squamate Reptiles. *Mol. Phylogenet. Evol.* **47**, 129–142 (2008).

113. N. R. Casewell, S. C. Wagstaff, R. A. Harrison, W. Wüster, Gene Tree Parsimony of Multilocus Snake Venom Protein Families Reveals Species Tree Conflict as a Result of Multiple Parallel Gene Loss. *Mol. Biol. Evol.* **28**, 1157–1172 (2011).

**Supplementary Acknowledgments**

The authors wish to thank the following individuals who very kindly helped with sampling for the phylogenetic dataset:

Bioken (Watamu, Kenya: Sanda Ashe, Anthony Childs, Royjan Taylor), Susen Baier, Patrick Barrière, Joe Beraducci, Phil Berry, Bill Branch, M. Brand, Donald G. Broadley, Ashok Captain, Tom Charlton, Laurent Chirio, F.W.P. Cotteril, Merel J. Cox, Liverpool School of Tropical Medicine (Edouard Crittenden, Paul D. Rowley, R. David G. Theakston), Jenny Daltry, Anslem de Silva, Latoxan (Yvon Doljansky, Franck Principaud), Craig Doria, California Academy of Sciences (Bob Drewes, Joe Slowinski), Damien Egan, Thomas Eimermacher, Johannes Els, Bryan G. Fry, Eli Greenbaum, Michael Griffin, Michel Guillot, Chris Hay, Hans-Werner Herrmann, Ivan Ineich, Andrew Jackson, Kate Jackson, Moshe Kahn, Trinh Xuan Kiem, Warren Klein, Lukas Kratochvil, Adam Leaché, Jonathan Leakey & Dena Crain, Thea Litschka-Koen, Mario Lutz, Anita Malhotra, Youssouph Mané, Bryan Maritz, Mark Marshall, Richard Mastenbroek, Peter Mirtschin, Zoltan Nagy, Deon Naude (Meserani), Mark O'Shea, Mike Perry, Tony Phelps, Jens B. Rasmussen, Radko Samek, Kate Sanders, Vishal Santra, Donald Schultz, Sterrin Smallbrugge, Martin Smit, Jeremie Tai-A-Pin, Zoltan Takacs, Bernard Thorens, Roger S. Thorpe, Jean-François Trape, Tanith Tyrr, Romilly van den Bergh, Craig van Rensburg, Mark Vandewalle, David A. Warrell, Zoological Society of London (Heather Hall, Terry March, Esther Wenman), Romulus Whitaker, Chris Wild, David Williams, Marcel Witberg, Addison Wynn, Luke Yeomans, Matthew Yuyek, London Zoo, Chai Koh Shin.
